# Rad54 separation of function mutation highlights unique roles during homologous recombination

**DOI:** 10.1101/2025.10.15.682645

**Authors:** Jingyi Hu, David Moraga, Amanda Xu, Lauren Peysakhova, J. Brooks Crickard

## Abstract

Homologous recombination (HR) is a DNA repair pathway that utilizes a template-based approach to repair double-strand breaks within the genome. Template utilization requires the exchange of individual DNA strands, which members of the RecA family of recombinases facilitate. Rad51 is a primary strand exchange factor in eukaryotes. During regular mitotic DNA repair, Rad51 is aided by the DNA translocase Rad54, which acts as a motor to remodel the template DNA and stabilize primary-strand exchange intermediates. The regulation of this activity remains incompletely understood. Here, we have identified a conserved site within the C-terminal region of Rad54. The mutation of this site creates a functional separation at early strand-exchange intermediates *in vivo*. Using this mutant protein, we identify a novel intermediate essential for stabilizing displacement loop (D-loop) structures. This precedes the removal of Rad51 and DNA extension. Based on our experiments, we hypothesize that this Rad54 mutant cannot stabilize Rad51-mediated strand-exchange intermediates because it cannot topologically isolate dsDNA regions. Identifying a mutant that disrupts this intermediate before Rad51 removal unifies existing models of Rad54-mediated D-loop formation and extension.

---

Homologous recombination (HR) is a DNA double-strand break repair (DSBR) pathway that uses a homologous template to prime DNA synthesis to repair breaks (1–5). In this pathway, ssDNA guides are formed by resection of the two ends of broken DNA (6–9). This DNA is known as the recipient DNA because it will receive information during the recombination reaction. Filaments of the RecA family of recombinases form on the recipient DNA (10–13). The filament guides a search of the genome for a matching DNA sequence (11,14–18). Once a suitable donor homology is identified, the recombinase filament initiates a strand exchange reaction that generates a three-stranded DNA joint known as a displacement loop (D-loop) (12,19,20). DNA polymerase can then initiate synthesis from the 3’ end of the recipient DNA and restore any lost information.

In the classic double-strand break repair pathway, resolution of DNA joints formed during HR can result in non-crossover outcomes (NCO) or crossover outcomes (CO). These outcomes are defined by the extent of genetic exchange between the donor and recipient, with CO outcomes resulting in greater exchange between the two DNA molecules (1,4). Resolution can also occur through synthesis-dependent strand annealing (SDSA). This pathway results in apparent NCO outcomes when the second end of the break is located and used to finish the repair. When the second end of the break is not located, SDSA can transition to Break-induced replication (BIR).

This outcome can be extremely mutagenic, leading to complex genomic rearrangements and mosaic repair (21–23).

In eukaryotes, the mitotic RecA homolog is known as Rad51 (24,25), and during the homology search and subsequent transition to DNA synthesis, Rad51 is assisted by the DNA motor protein Rad54 (26–32). The basic biochemical properties of Rad54 are that it hydrolyzes ATP to physically move along double-stranded DNA by tracking the minor groove in the 3’-5’ direction (33). Multiple copies of Rad54 can function as single units, resulting in high processivity levels (27,29,32). The primary role of Rad54 is to act as an accessory cofactor for Rad51, and interactions between these proteins increase the motor output of Rad54 by 3 to 5-fold (29,34,35). The activities of Rad54 include the regulation of Rad51 at stalled or collapsed DNA replication forks (36), the removal of excess Rad51 that is pathogenically bound to dsDNA (37,38), the remodeling of nucleosomes (39–41), driving branch migration (35,42–45), collaboration with Rad51 to catalyze the formation of D-loops (19,29,32,46–48), and to grant accessibility of the 3’ end of the recipient DNA to DNA polymerase (47,49).

*In vivo*, Rad54 can promote Rad51 recombination activity through multiple mechanisms. One of these involves the removal of Rad51 from dsDNA and its stabilization on ssDNA (28,37,50–53), which aids in the maturation of recombinase filaments. Evidence for this stems from work involving the Rad51 Walker A mutant, Rad51K191R, which is deficient in filament formation *in vivo* (54). The overexpression of Rad54 can suppress the defect in this mutant, but only in the presence of Rad55/57 (55). Rad55/57 are required for efficient Rad51 filament formation and counteracting the effects of the Srs2 helicase (56–60), whose primary function during HR is to remove Rad51 from ssDNA (61–66). During Rad51 filament maturation, Rad54 can act by increasing the available pools of Rad51 and by physically protecting Rad51 filaments from Srs2 activity (53). Srs2 and Rad54 are synthetically lethal (67) and appear to compete during repair. This competition likely occurs before the alignment of donor and recipient DNA and can occur before initial invasion or during reinvasion events during BIR-based repair (22,23).

This model has been extended to support the hypothesis that the removal of Rad51 from newly paired dsDNA at strand-exchange intermediates is how Rad54 stabilizes the three-stranded D-loop structures (47). This hypothesis explains the generation of an accessible 3’ end for further DNA extension, promoting repair and intermediate stability. A key feature of this model is that there is no functional separation in Rad54 activity. A contrasting model posits that Rad54’s activities are distinct functions, and that its role in forming D-loops does not require the removal of Rad51 from dsDNA (46). The basis for this hypothesis is that Rad54 generates underwound DNA during translocation, and that negatively supercoiled DNA is a more efficient substrate for recombination (29,46,68). Whether D-loop stabilization and Rad51 removal from dsDNA are distinct or overlapping functions remains unclear *in vivo*.

Here, we investigated potential phosphorylation sites on the Rad54 protein and identified a specific residue in the C-terminal region that modulated activity. While this residue could not be confirmed as phosphorylated, it yielded a separation-of-function mutant. We show that the substitution of Aspartic acid for two Serine residues disrupts a bridging contact between the two RecA lobes of Rad54. Disrupting this contact caused severe defects in D-loop formation *in vivo* but allowed for D-loop extension and the removal of Rad51 from dsDNA. Our data suggest the identification of a novel physiological intermediate during D-loop formation, indicating that this step likely stabilizes the early D-loop and potentially later recombination intermediates. We hypothesize that this is due to Rad54’s ability to clamp dsDNA during strand exchange, thereby inducing force-mediated remodeling of the donor DNA. The removal of Rad51 follows this. By using this mutation, our study shows that these steps are mechanistically distinct, reconciling and unifying two existing models of Rad54-mediated D-loop formation.

## Results

We identified potential phosphorylation sites in *Saccharomyces. cerevisiae* Rad54 using the SuperPhos database (69). Seven sites were selected, and these residues were mutated to Aspartic acid or Alanine for Serine, and to Glutamic acid or Alanine for Threonine. We performed complementation assays in *rad54* strains using methyl methanesulfonate (MMS) as a DNA-damaging agent (**Figure 1A**). Most of the mutated residues successfully complemented the MMS sensitivity phenotype. The exception was that the substitution of Aspartic acid for Serine at residue 816, which failed to fully complement the MMS sensitivity of the *rad54* strain (**Figure 1B**). We further inspected the S816 site and found that the adjacent residue was also a Serine (**Figure 1C**). We speculated that mutating both residues may result in a more severe phenotype, so we generated *rad54S816A, S817A,* and *rad54S816D, S817D* substitutions and tested them for complementation. We observed an enhanced phenotype in the *rad54S816D, S817D* strain (**Extended View Figure 1A**). There was no observable phenotype when only the S817 was changed to an Aspartic acid residue (**Extended View Figure 1A**). This implies that the S816 site is the most critical amino acid.

**Figure 1:**
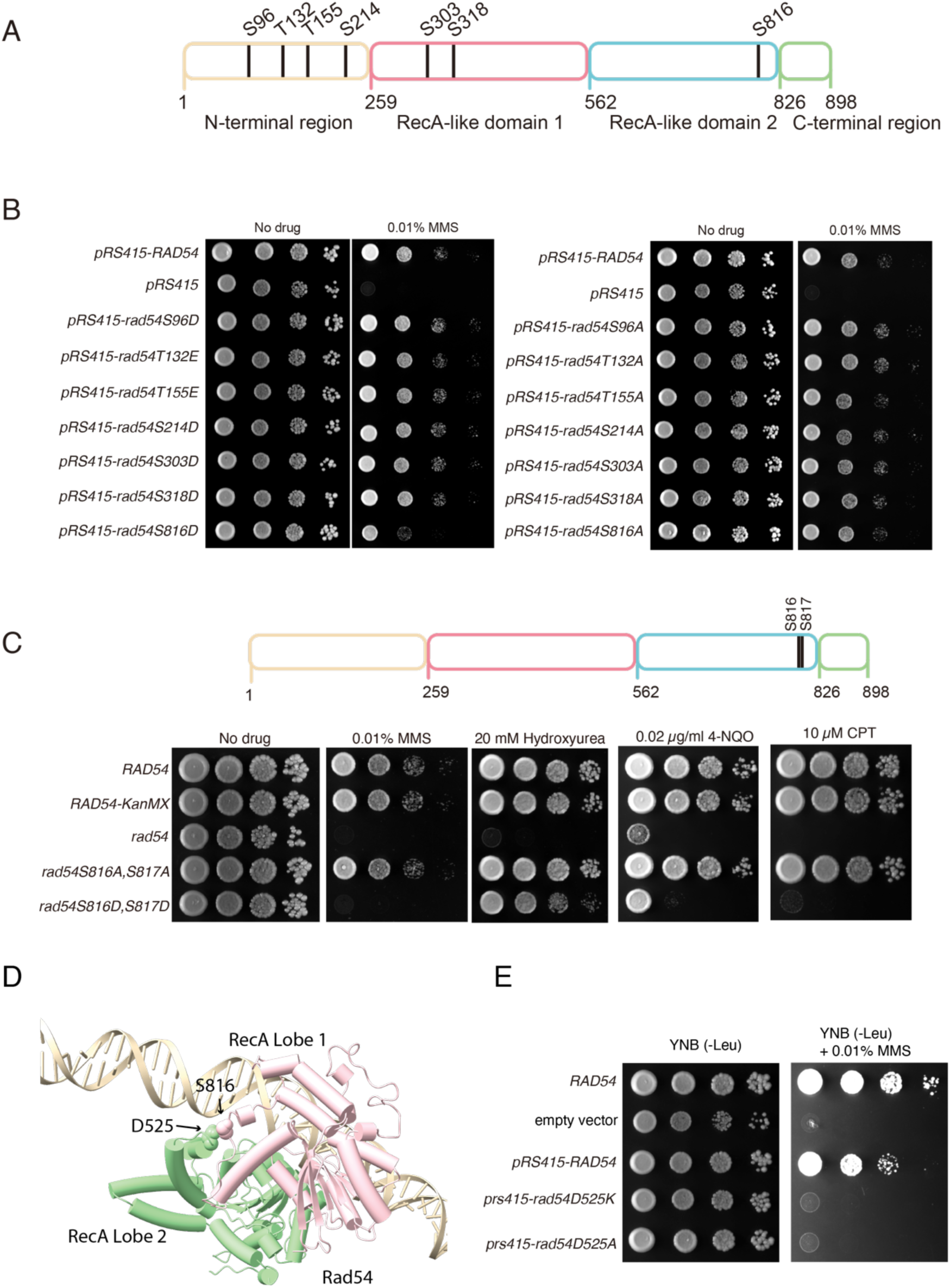
Screen of potentially phosphorylated residues in Rad54 **(A).** Schematic of residues in Rad54 that were identified as phosphorylated in the SuperPhos database. **(B).** Yeast complementation spot assay to monitor the effect of phosphomimic mutants (Left) and alanine mutants (Right) of Rad54 at 0.01% MMS. **(C).** Serial dilution spot assay to monitor the impact of *rad54S816A, S817A*, and *rad54S816D, S817D* on sensitivity to 0.01% MMS, 20 mM Hydroxyurea (HU), 0.02 µg/ml 4-NQO, and 10 µM CPT. **(D).** AlphaFold generated model of Rad54 bound to dsDNA. The two RecA lobes are color coded, and residues D525 and S816 are shown as spheres. **(E).** Serial dilution spot assay showing sensitivity of *RAD54,* empty vector, *pRS415-RAD54, pRS415-rad54D525K*, and *pRS415-rad54D525A* to 0.01% MMS,

Under the MMS conditions used for the initial screen, *rad54S816D, S817D* were as severe as the *rad54* strains. Therefore, we tested whether this mutation could be expressed at similar levels to RAD54 and whether it could still form foci in response to MMS treatment. We monitored *RAD54-GFP*, *rad54S816A, S817A-GFP,* and *rad54S816D, S817D-GFP* for expression with and without MMS (**Extended View Figure 2A**). All three of these proteins were expressed at similar levels with and without MMS and formed Rad54 foci in response to MMS treatment (**Extended View 2ABC**). From this, we concluded that there was no defect in the expression or stability of these Rad54 mutants.

**Figure 2:**
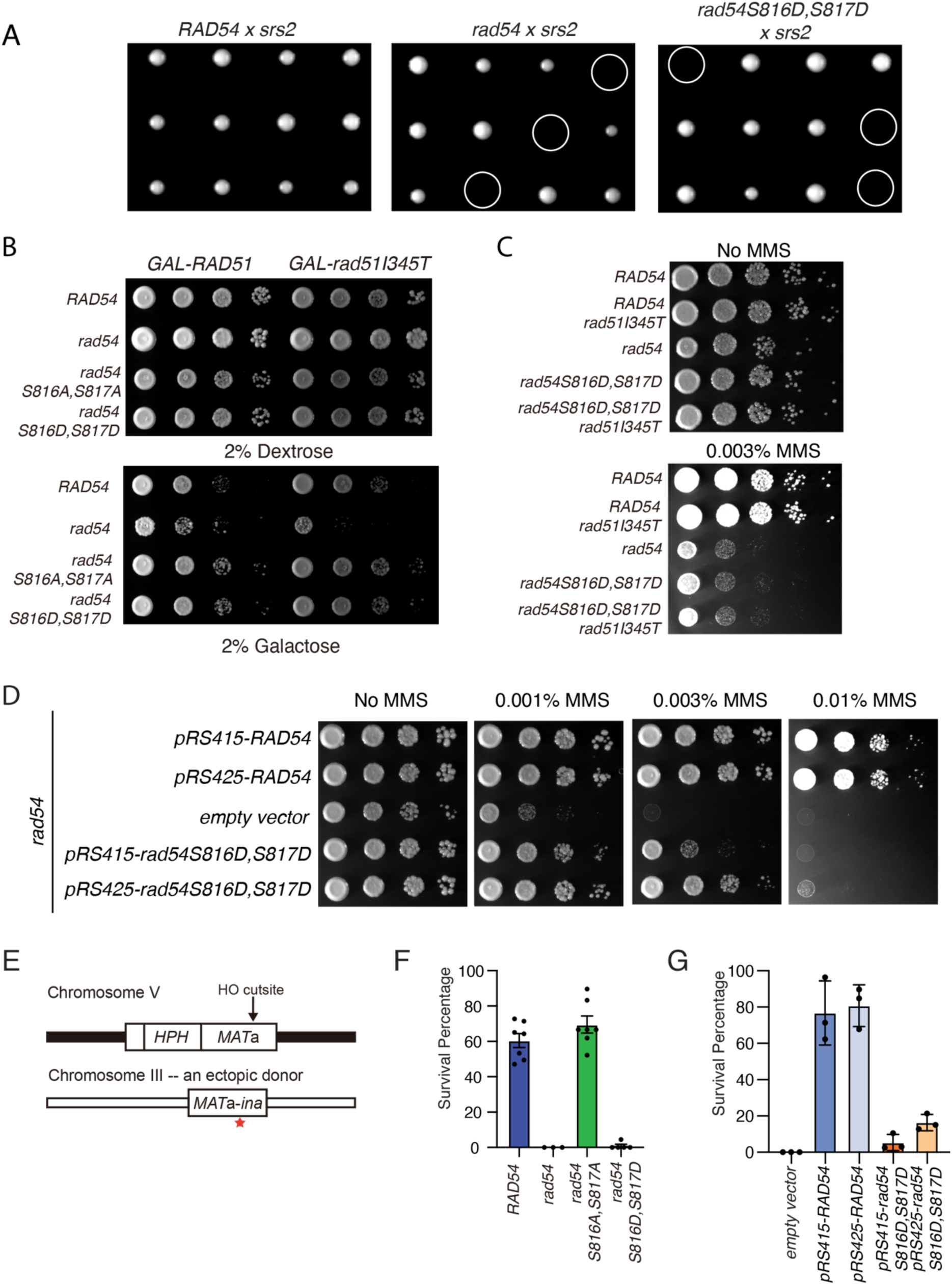
Mutations in Rad54 are separation-of-function mutants **(A).** Representative tetrad analysis for *RAD54 x srs2*, *rad54 x srs2,* and *rad54S816D, S817D x srs2*. Each row is separated from one tetrad. White circles highlight synthetic lethal combinations. **(B).** Serial dilution spot assay to determine the effect of GAL-*RAD51* and GAL-*rad51I345T* over-expression on *RAD54, rad54, rad54S816A, S817A,* and *rad54S816D, S817D* strains. **(C).** Serial dilution spot assay to test the MMS sensitivity of *RAD54, RAD54 rad54I345T, rad54, rad54 S816D, S817D*, and *rad54 S816D, S817D rad51I345T* **(D).** Serial dilution spot assay for rad54 complemented with *pRS415-RAD54, pRS425-RAD54*, empty vector, *pRS415-rad54S816D, S817D*, and *pRS425-rad54S816D, S817D*. Strains were tested with no MMS, 0.001%, 0.003%, and 0.01% MMS. **(E).** Schematic for the experiment used to measure the repair of a double strand break at an ectopic site. **(F).** Graph representing the colony survival rate for *RAD54, rad54, rad54S816A, S817A, rad54S816D, S817D.* The bars represent the mean, and the error bars represent the standard error measurement of at least three independent experiments. **(G).** Bar graph representing analysis of *RAD54* overexpression on the complementation of repair of an ectopic double-strand break. All strains are *rad54* with the empty vector, *pRS415-RAD54, pRS425-RAD54, pRS415-rad54S816D,S817D,* and *pRS425-rad54S816D,S817D*. The bar represents the mean, and the error bars represent the standard error measurement of three independent experiments.

We evaluated the sequence around the S816 site for a potential kinase consensus sequence. The surrounding sequence appeared to be a degenerate polo-like kinase site (**Extended View figure 3A**) (70). In human RAD54L, the site is a complete pololike kinase consensus sequence. The yeast polo-like kinase is Cdc5, which is implicated in the resolution of recombination intermediates (71,72). We made several attempts to confirm that this site was phosphorylated but were unsuccessful. Therefore, we can only conclude that the previously established phosphoproteomics has indicated this as a phosphorylation site. We next inquired whether this residue was structurally important by performing multiple sequence alignments of 1200 Rad54 sequences from eukaryotes (73) and found that S816 was conserved in 94.7% of sequences. One of the few exceptions was Rad54 from Dictyostelium, in which the Serine was replaced with an Aspartic acid residue (**Extended View Figure 3B**). The natural replacement of Serine by Aspartic acid suggested that *rad54S816D, S817D* may retain partial function.

**Figure 3:**
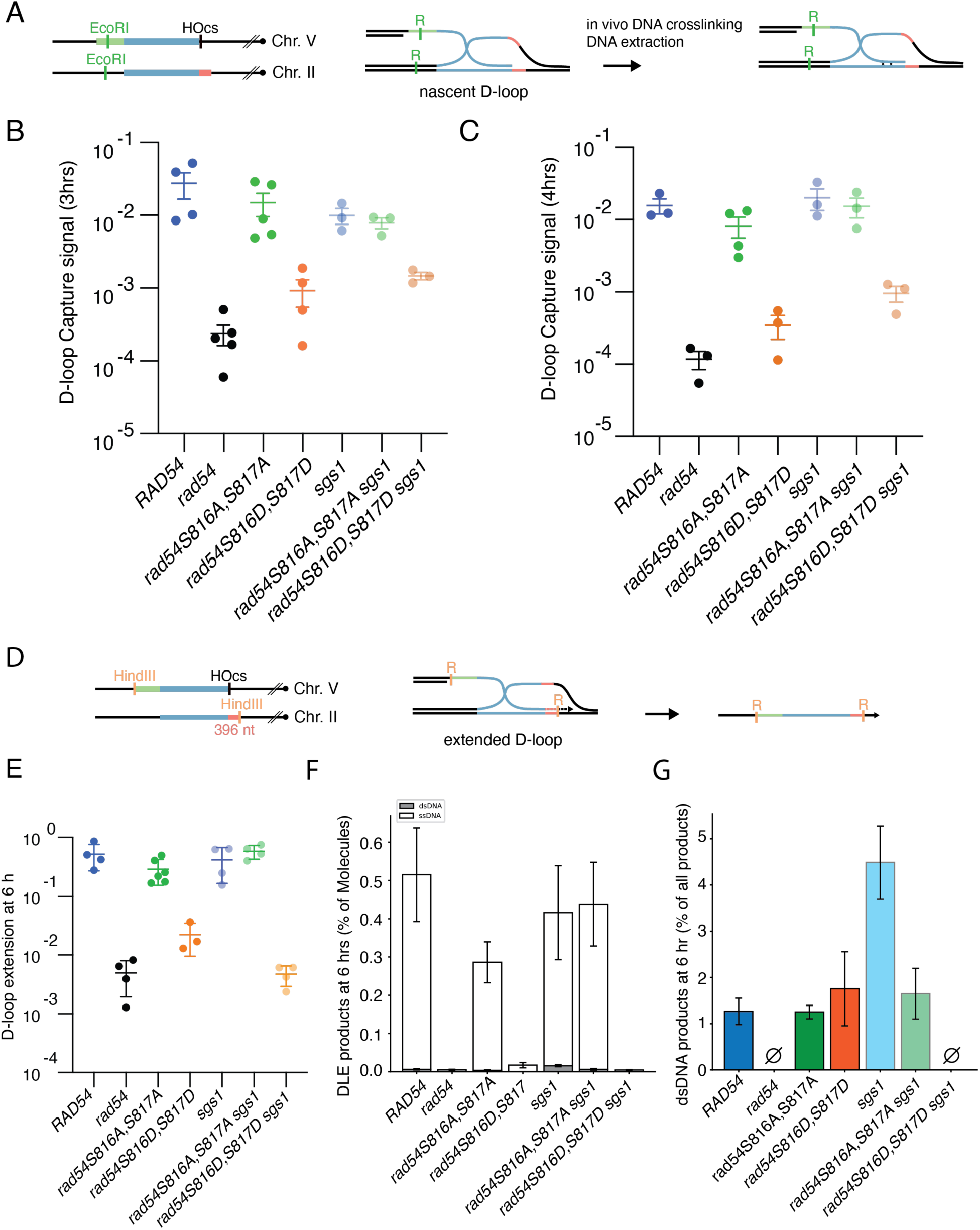
Mutations in *RAD54* impact recombination intermediates **(A).** Schematic diagram illustrating the D-loop capture assay to trap the formation of nascent D-loops during recombination. **(B).** Graph representing D-loop capture efficiency at 3 hours post-break induction for *RAD54*, *rad54*, *rad54S816A, S817A*, *rad54S816D, S817D*, sgs1, *rad54 S816A, S817A sgs1*, *rad54S816D, S817D sgs1.* The bar represents the mean, and the error bars represent the standard error measurement for at least three independent experiments. **(C).** Graph representing D-loop capture efficiency at 4 hours post-break induction for *RAD54*, *rad54*, *rad54 S816A, S817A*, *rad54S816D, S817D*, sgs1, *rad54S816A, S817A sgs1*, *rad54S816D, S817D sgs1.* The bar represents the mean, and the error bars represent the standard error measurement for at least three independent experiments. **(D).** Schematic diagram illustrating the assay to monitor D-loop extension. **(E).** Graph representing D-loop extension at 6 hours for *RAD54*, *rad54*, *rad54 S816A, S817A*, *rad54S816D, S817D*, sgs1, *rad54S816A, S817A sgs1*, *rad54S816D, S817D sgs1*. The bar represents the mean, and the error bars represent the standard error measurement for at least three independent experiments. **(F).** Extension products at 6 hours: single-stranded (ssDNA; DLE signal when both hybrid oligos were added) and double-stranded (dsDNA; DLE signal when no hybrid oligos were added) for *RAD54*, *rad54*, *rad54 S816A, S817A*, *rad54S816D, S817D*, sgs1, *rad54S816A, S817A sgs1*, *rad54S816D, S817D sgs1*. The error bars represent the standard error measurement of at least three independent experiments. **(G).** The percentage of dsDNA among total extension products for *RAD54*, *rad54*, *rad54 S816A, S817A*, *rad54S816D, S817D*, sgs1, *rad54S816A, S817A sgs1*, *rad54S816D, S817D sgs1*. The error bars represent the standard error measurement for at least three independent experiments.

To determine whether *rad54S816D, S817D* retained partial function, we measured its ability to complement different DNA-damaging agents. MMS is an alkylating agent primarily repaired by the base excision repair pathway (74). However, at sufficiently high concentrations, HR is required to repair DNA breaks caused by stalled replication forks or transcription stress. Because different DNA-damaging agents utilize distinct repair pathways, we evaluated whether *rad54S816D, S817D* was sensitive to other types of damaging reagents. Hydroxyurea (HU) is a drug that depletes nucleotide pools and acts as a damaging agent by inducing replication stress (75). We observed that *rad54* strains are unable to grow in the presence of 20 mM HU (**Figure 1C**). In contrast, we observed only a minor growth defect for *rad54S816D, S817D,* suggesting that this protein retains some function and is relatively unaffected by HU.

Camptothecin (CPT) is a topoisomerase inhibitor that creates protein-DNA adducts (76). These adducts are primarily repaired via cross-link repair pathways. However, at sufficiently high concentration, these adducts cause transcription and replication stress that requires HR for repair. Under CPT treatment conditions, we observed an odd effect. Experiments performed with BY4741 strains obtained from the Dharmacon collection showed that rad54 strains were completely *deficient*. In contrast, *rad54S816D, S817D* showed only minor defects (**Extended View Figure 4**). This was not the case using a model strain from the laboratory, which showed severe CPT sensitivity in the *rad54S816D, S817D* mutant (**Figure 1C**). We are uncertain why this difference occurred, but it is likely due to differences in the genetic backgrounds of these strains, which may make the lab model strain more dependent on HR to repair CPT-induced lesions. Finally, we tested sensitivity to 4-Nitroquinoline Oxide (4-NQO). This drug is a UV-mimetic chemical that forms base adducts (77). At sufficiently high concentrations, it can lead to transcription- or replication-induced breaks. Treatment with 4-NQO resulted in sensitivity in the *rad54S816D, S817D* strains. From this, we conclude that *rad54S816D, S817D* retain partial function under some DNA-damaging conditions, depending on the type of damage.

**Figure 4:**
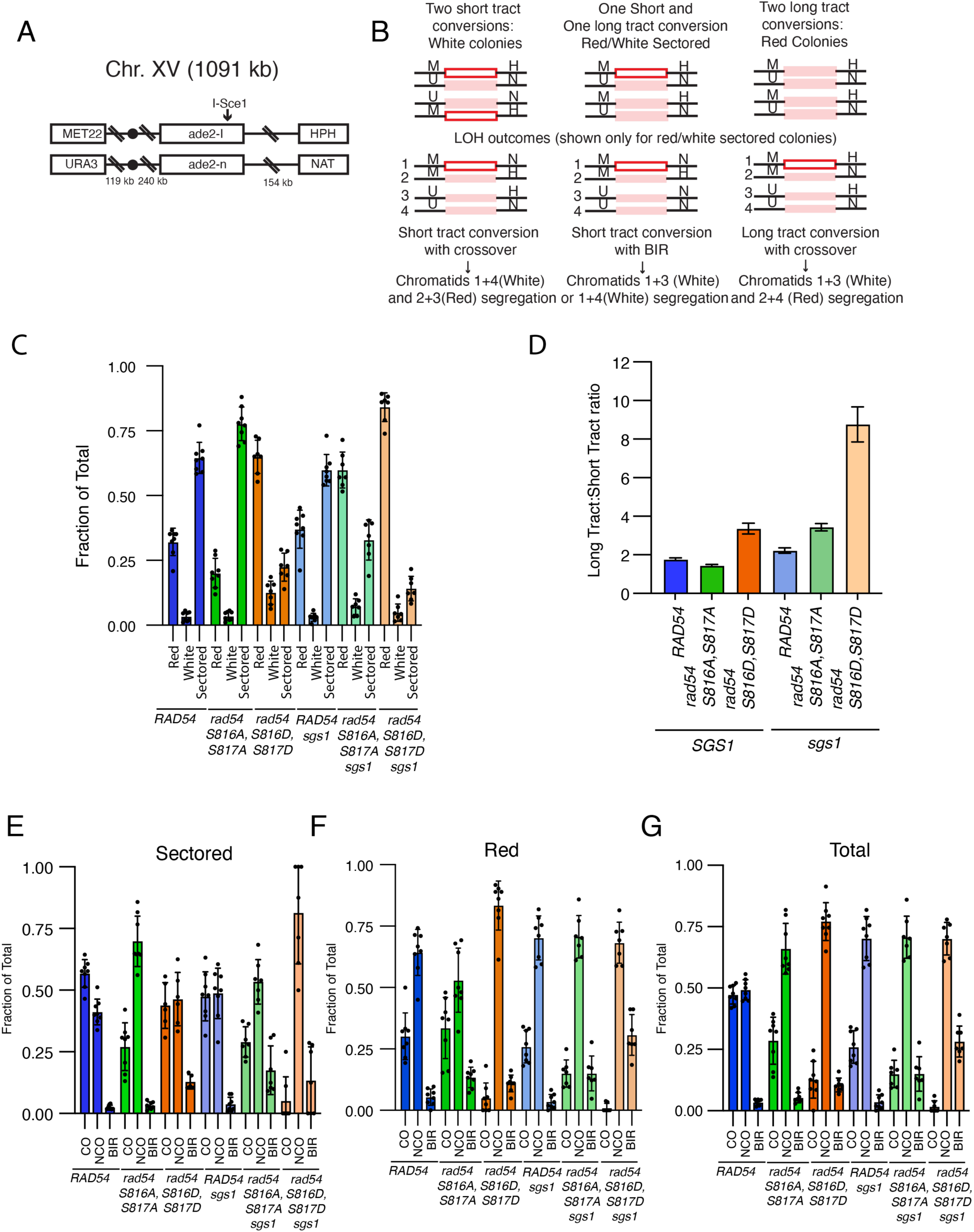
Impact of *RAD54* mutations on allelic recombination **(A).** Schematic diagram illustrating the DNA reporter used to analyze the effect of Rad54 mutation on allelic recombination. **(B).** Schematic diagram illustrating the potential gene conversion outcomes and HR based repair pathways during allelic recombination. **(C).** A bar graph representing the gene conversion outcomes for *RAD54*, *rad54S816A, S817A*, and *rad54S816D, S817D*. The bars represent the mean of the data, and the error bars represent the standard deviation. The data are the result of at least 6 independent experiments. **(D).** Graph illustrating the ratio of long tract to short tract gene conversion for *RAD54, rad54S816A, S817A, and rad54S816D, S817D*. The error bars represent the standard error of the measurement. **(E).** Bar graph representing the recombination outcomes for sectored colonies for *RAD54, rad54S816A, S817A, and rad54S816D, S817D.* The bars represent the mean, and the error bars represent the standard deviation of the data. The data are the results of at least 6 independent experiments. **(F).** Bar graph representing the recombination outcomes for red colonies for *RAD54, rad54S816A, S817A, and rad54S816D, S817D.* The bars represent the mean, and the error bars represent the standard deviation of the data. The data are the results of at least 6 independent experiments. **(G).** Bar graph representing the total recombination outcomes for *RAD54, rad54S816A, S817A, and rad54S816D, S817D.* The bars represent the mean, and the error bars represent the standard deviation of the data. The data are the results of at least 6 independent experiments.

To better understand the defects associated with the substitution of Serine 816 with Aspartic acid, we predicted the structure of *S. cerevisiae* Rad54 (**Figure 1D and Extended View Figure 5A**) and *H. sapiens* RAD54 bound to 60 bp of dsDNA (**Extended View Figure 5B**) using AlphaFold3 (78). The S816 residue in *Sc*Rad54 and S657 in *H.s.*RAD54 formed a bridging interaction by contacting a conserved (**92.5%)** Aspartic acid residue (D525 in *S.c.*Rad54 and D366 in *H.s.*RAD54) (**Figure 1D and Extended View Figure 5**CD). This interaction was not observed in the *Danio. rerio* crystal structure (79), likely because it lacks double-stranded DNA (dsDNA). We also evaluated the data from the AlphaFold repository, which predicts the pathogenicity of specific amino acid substitutions at a given site for human proteins (**Extended View Figure 5E**) (80). This analysis indicated that substitutions at S657 (equivalent to S816 in *S.c.* Rad54), L658 (equivalent to S817 in *S.c.* Rad54), and D366 (equivalent to D525 in S.c.Rad54) could potentially be pathogenic, depending on the specific amino acid substitution.

**Figure 5:**
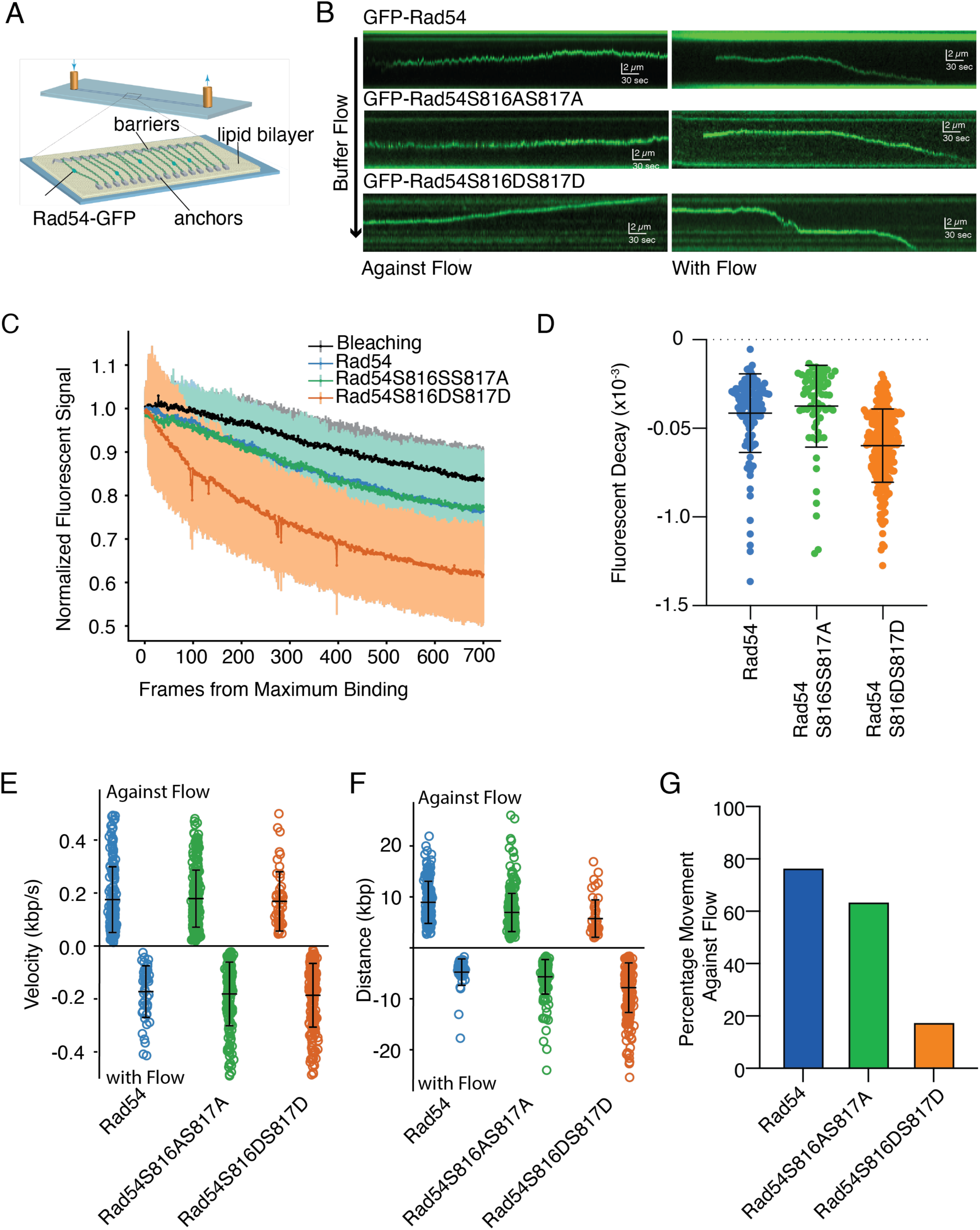
Rad54S816D/S816D has reduced affinity for dsDNA **(A).** Schematic diagram illustrating DNA curtains experiments to test the activity of Rad54 on dsDNA. **(B).** Representative kymographs for GFP-Rad54, GFP-Rad54 S816A/S817A, and GFP-Rad54 S816D/S817D moving against (Left) and with (Right) buffer flow. **(C).** Mean fluorescent decay rate for Rad54, Rad54 S816A/S817A, and Rad54 S816D/S817D. The black line gives the photobleaching rate. The shade represents the standard deviation of the data **(D).** Dot plot with the individual fluorescence decay rates for Rad54 (N=117), Rad54 S816/AS817A (N=68), and Rad54 S816D/S817D (N=232). The bar represents the mean, and the error bars represent the standard deviation of the data. **(E).** Graph representing the velocity of translocation in kbp/s for Rad54 (N=216), Rad54 S816A/S817A (N=446), and Rad54 S816D/S817D (N=271). The molecules that moved against the flow are above the X-axis, and the molecules that moved with the flow are below the X-axis. The bar represents the mean, and the error bars represent the standard deviation of the data. **(F).** Graph representing the distance moved for Rad54 (N=216), Rad54 S816A/S817A (N=446), and Rad54 S816D/S817D (N=271). The molecules that moved against the flow are above the X-axis, and the molecules that moved with the flow are below the X-axis. The bar represents the mean, and the error bars represent the standard deviation of the data. **(G).** Bar graph representing the percentage of Rad54 (166/216), Rad54 S816A/S817A (285/446), and Rad54 S816D/S817D (48/271) that move against the buffer flow.

To further validate these observations, we generated plasmids encoding *rad54D525K* and *rad54D525A* and tested their ability to restore the sensitivity of *rad54* strains to DNA-damaging agents. We found that both *rad54D525K* and *rad54D525A* failed to complement MMS sensitivity (**Figure 1E**). Like the *rad54S816D, S817D* substitution, there was a failure to fully complement CPT and 4-NQO sensitivity, although the phenotype was not as severe as the *rad54S816D, S817D* mutant (**Extended View Figure 6A**). We also observed that this substitution complemented HU phenotypes and therefore retained partial function (**Extended View Figure 6A**). From this analysis, we conclude that the likely defect observed in the *rad54S816D, S817D* mutant is due to a disruption between the two RecA lobes of Rad54.

**Figure 6:**
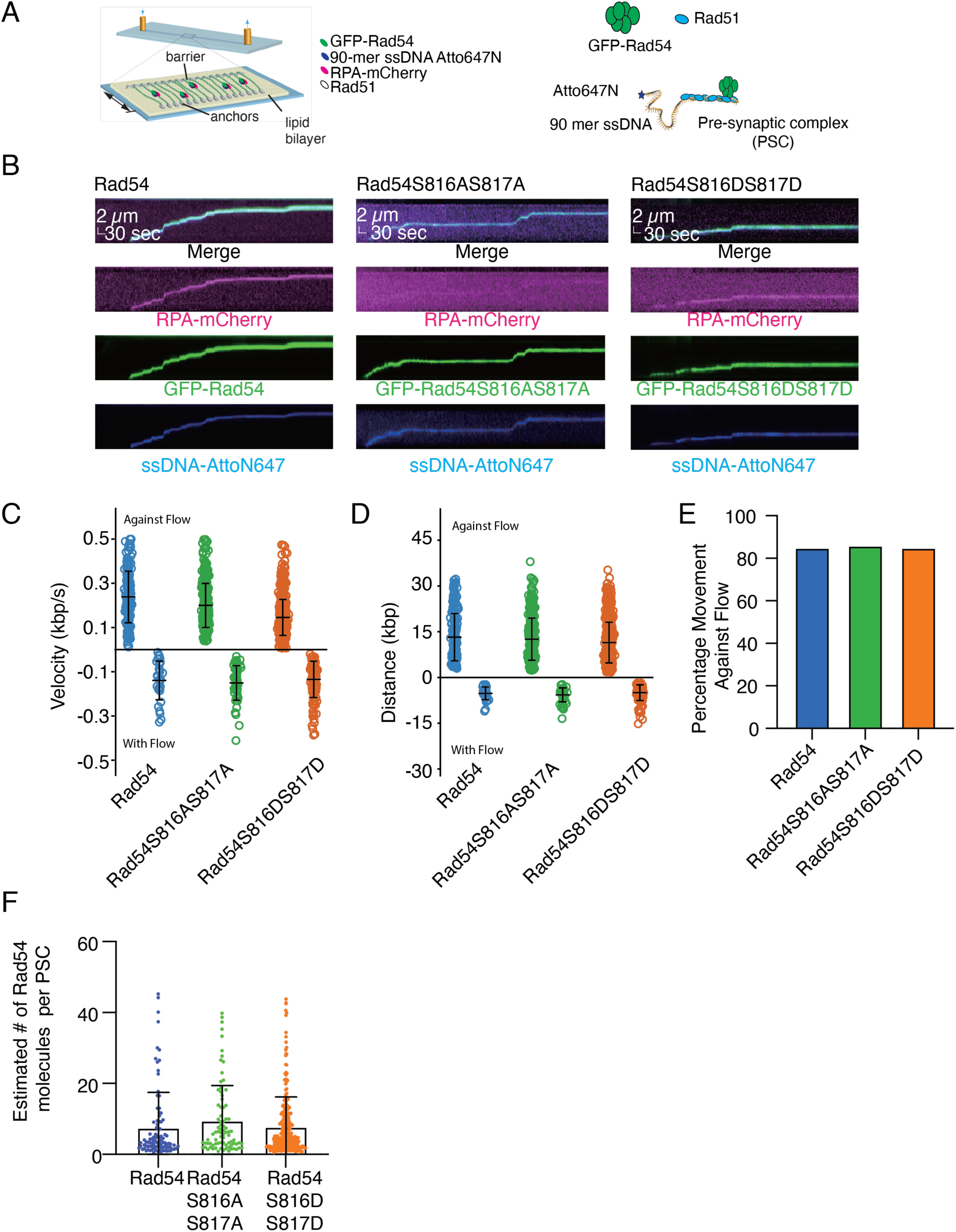
Weak general defects in Rad54 S816D/S817D in the PSC **(A).** Cartoon diagram illustrating the use of DNA curtains to perform experiments to monitor the activity of the presynaptic complex (PSC) on dsDNA. **(B).** Representative kymographs for PSCs, PSCs with Rad54 S816A/S817A, or PSCs with Rad54 S816D/S817D. Shown are merged images of RPA-mCherry, GFP-Rad54, and Atto647-90-mer ssDNA. **(C).** Measured translocation velocities for the PSC with Rad54 (N=215), PSC with Rad54 S816A/S817A (N=378), and PSC with Rad54 S816D/S817D (N=835). The PSCs that moved against the flow are above the X-axis, and the PSC values that moved with the flow are below the X-axis. The bars represent the mean, and the error bars represent the standard deviation of the data. **(D).** Measured translocation distances for the PSC with Rad54 (N=215), PSC with Rad54 S816A/S817A (N=378), and PSC with Rad54 S816D/S817D (N=835). The PSCs that moved against the flow are above the X-axis, and the PSC values that moved with the flow are below the X-axis. The bars represent the mean, and the error bars represent the standard deviation of the data. **(E).** Graph representing the percentage of PSC with Rad54 (183/215), PSC with Rad54 S816A/S817A (326/378), and PSC with Rad54 S816DS817D (710/835) that moved against the buffer flow. **(F).** Graph representing the estimated number of Rad54 molecules per PSC with Rad54 (N=96), Rad54S816A/S817A (N=82), and Rad54S816D/S817D (N=271). The dot represents the mean, and the error bars represent the standard deviation of the experiment.

## *rad54 S816D,S817D* can perform presynaptic *RAD54* functions

The presynaptic phase of HR based repair is characterized by the growth and remodeling of Rad51 filaments on ssDNA. The UvrD helicase Srs2 plays an important role in this step of recombination, modulating the length of Rad51 filaments (62,63,65,66). This role appears to be required for proper Rad51 filament maturation, which is at least needed for ectopic recombination (65). Under these conditions, the activity is pro-recombinogenic. While the pre-synaptic phase occurs widely on ssDNA generated during resection, it can also occur when long stretches of ssDNA are generated during break-induced replication (BIR) (22,81–83). Under these conditions, Srs2 acts as an anti-recombinase, reducing toxic intermediates that arise from potential reinvasion. At stalled replication forks, Srs2 can also act as an anti-recombinase, preventing HR and promoting translesion synthesis (65,84). Rad54 competes with Srs2 by stabilizing Rad51 filaments and preventing promiscuous binding of Rad51 to dsDNA, thereby increasing the active pool of Rad51. Based on this Rad54 activity, if *rad54S816D, S817D* were only defective in the presynaptic phase of HR, we might expect that this mutant would no longer be synthetically lethal with *SRS2,* would be unable to remove Rad51 from dsDNA, and the phenotype would be reversible with overexpression.

*RAD54* and *SRS2* are synthetically lethal. Typically, synthetic lethality occurs when genes are required for parallel or compensatory pathways. Divergence of these two pathways may occur during a modified presynaptic phase of HR at stalled replication forks, where HR (RAD54) competes with translesion synthesis (SRS2) (84). In this scenario, Rad54 likely antagonizes Srs2 to promote HR by preventing Rad51 from being removed. We reasoned that if *rad54 S816D, S817D* were defective in the presynaptic phase, it would not be synthetically lethal with *SRS2,* as this would allow the use of the SRS2-favored pathway. We performed tetrad analysis to measure synthetic lethality. We compared *RAD54 srs2*, *rad54 srs2*, and *rad54 S816D, S817D srs2*. The tetrad dissection pattern showed that the *rad54 S816D, S817D* is synthetically lethal with *srs2* (**Figure 2A**). This was not due to defects in meiosis because the spores followed a 1:1:4 (Parental Ditype (DP), Non-Parental Ditype (NPD), Tetra Type (TT)) segregation pattern.

Rad54 can remove Rad51 bound to dsDNA (37,38,51,85). This activity may participate in the recovery of stalled DNA replication forks (36), the stabilization of the nascent D-loop (47), or increase the active pools of Rad51. Overexpression of the recombinase Rad51 can lead to pathological binding of Rad51 to double-stranded DNA (37,38). When Rad51 is overexpressed in a r*ad54* strain, there is a minor growth phenotype. We reasoned that if *rad54 S816D, S817D* retained this function, then it would suppress this phenotype. We found complementation of this phenotype by both *rad54S816A, S817A*, as well as *rad54S816D, S817D* (**Figure 2B).** This result indicates that these Rad54 mutants can remove Rad51 from dsDNA. To further challenge the mutants, we also performed this experiment with *rad51I345T* overexpression. This mutant form of Rad51 binds to dsDNA more efficiently (57) and should be more difficult to remove. However, overexpression of this Rad51 variant did not significantly impact the *rad54* phenotype or prevent complementation by the Rad54 mutants, compared with overexpression of wild-type Rad51 (**Figure 2B**). From this, we conclude that *rad54 S816D, S817D* is competent in removing Rad51 from dsDNA. *rad51I345T* is a Rad51 variant that also produces longer Rad51 filaments and suppresses the *SRS2* deletion phenotype. If *rad54S816D, S817D* were defective in presynaptic activity, we may expect *rad54I345T* to suppress the MMS sensitivity observed in these strains. However, we observed no suppression when we tested this strain (**Figure 2C**).

The activity of the presynaptic phase is strongly controlled by concentration due to stabilization of Rad51 filaments. As a result, Rad54 overexpression may lead to increased HR. If this were the primary defect in the *rad54S816D, S817D* mutant, then we would expect overexpression of the protein to rescue the DNA damage phenotype. Therefore, we expressed Rad54 from either a centromeric plasmid or a 2µ plasmid. The 2 µ plasmid is expected to result in 10- to 15-fold overexpression of the protein. When we tested the overexpression strains for complementation of the *rad54* MMS phenotype. We observed a decrease in sensitivity in cells expressing *rad54S816D, S817D* from the 2 µ plasmid (**Figure 2D**). However, this complementation was incomplete and did not fully restore *RAD54* resistance levels (**Figure 2D**). These data suggest that *rad54S816D, S817D* support presynaptic Rad54 activity but are defective later in the HR pathway.

## *rad54 S816D, S817D* is defective in stable D-loop formation

The ability of the *rad54S816D, S817D* variant to function before displacement loop formation, during the presynaptic phase of HR, suggests that it is a separation-of-function mutant. To fully address the stage at which Rad54 activity is disrupted, we evaluated whether there was a defect in repairing a single double-strand DNA break from an ectopic donor. We used a reporter assay with a MATa locus containing an HO endonuclease cleavage site on chromosome V (**Figure 2E**) (86). A homologous non-cleavable MATa is located on chromosome III (**Figure 2E**). Upon induction with galactose, the DNA is broken, and repair occurs using the ectopic homologous site. Cells will only survive if they can repair the break. We measured the survival percentage after induction of a break. In wildtype (WT) strains, the survival frequency was 60-65% (**Figure 2F**). This was also the case with the *Rad54S816A, S817A* strains (**Figure 2F**). In the *rad54* strain, there is no survival. The *rad54S816D, S817D* strains exhibited only ∼1% survival (**Figure 2F**). We further tested whether *rad54D525A* or *rad54D525K* could complement the DSB repair phenotype. With these mutants, the survival rate was approximately 25% (**Extended View Figure 6C**).

Finally, we tested whether overexpression of *rad54S816D, S817D* could complement the repair phenotype. When *rad54S816D, S817D* were expressed from a centromeric plasmid, only 3% of the cells survived, comparable to the gene replacement strains. Expression from the two µ plasmid resulted in ∼ 12% survival (**Figure 2G**). While this was significantly better than survival under other conditions, it was not as high as the 75-85% survival observed for RAD54 overexpression. From these experiments, we conclude that *rad54S816D, S817D* is defective in double-strand break repair.

To identify the repair intermediate at which the *rad54S816D, S817D* strain is defective, we used a previously established D-loop capture assay (14,19,87,88). This assay also uses an ectopic repair site within the genome. After induction of a break using the HO endonuclease, the cells are cross-linked with psoralen, which can trap the initial three-strand intermediates, known as nascent D-loops. The efficiency of D-loop capture is then quantified by proximity ligation and qPCR (**Figure 3A and Extended View Figure 7AB**). We monitored D-loop capture at 3 and 4 hours, and, as expected, the *rad54* strains showed 100-fold lower signals than the WT. This value was also the same as that obtained from qPCR performed without the oligo to restore restriction enzyme sites, indicating this is the background (**Extended View Figure 7C**). The *rad54S816A, S817A* were slightly lower than WT. However, this difference was not significant(**Figure 3B**). The *rad54S816D, S817D* strains were 34-and 47-fold lower than WT at 3 and 4 hours, respectively (**Figure 3B and Extended View Figure 7CDEG**), indicating a severe defect in D-loop capture.

**Figure 7:**
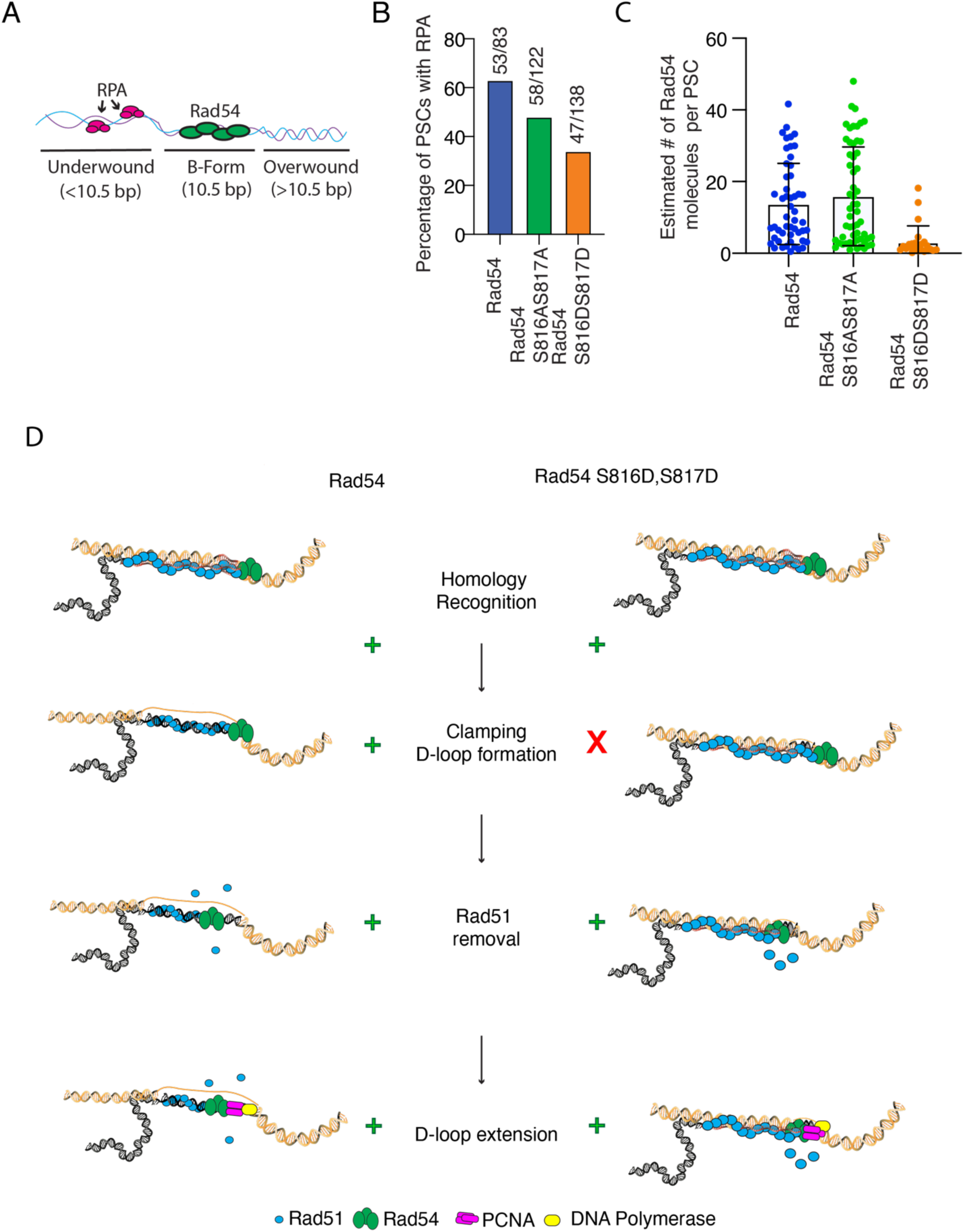
Disruption in Rad54 clamping reduced D-loop formation **(A).** Cartoon diagram illustrating the recruitment of RPA to regions of underwound DNA generated by the PSC. **(B).** Graph representing the percentage of PSCs (53/83), PSCs with Rad54 S816AS817A (58/122), and PSCs with Rad54 S816DS817D (47/138) that have RPA associated with them. **(C).** Graph representing the estimated number of GFP-Rad54 molecules bound to the PSC after a 5-minute flow-off incubation for Rad54 (N=51), Rad54S816A/S817A (N=56), and Rad54S816D/S817D (N=24). The bar represents the mean, and the error bar represents the standard deviation of the data. **(D).** Model illustrating the role of Rad54 during D-loop formation. On the left is the WT case, and on the right is the Rad54 S816D/S817D.

We next asked whether the *rad54S816D, S817D* mutations disrupted D-loop formation or were associated with an elevated level of nascent D-loop reversal. Sgs1 is a helicase implicated in the reversal of D-loops (19,87,89). We reasoned that if *rad54S816D,S817D* were subjected to a high level of D-loop reversal, then deletion of *SGS1* might suppress most of the observed defects in D-loop capture. However, if the increase were small, then it would imply that most of the defect is due to the initial capture. An increase in D-loop capture was observed in the *rad54S816D, S817D sgs1* double mutant strain. However, this was still 20- and 12-fold lower than WT at 3 and 4 hours, respectively (**Figure 3BC and Extended View Figure 7C**). The increase observed in the *rad54S816D,* S817D *sgs1* double mutants translates into 1.4- and 4-fold increases in capture efficiency compared with the *rad54S816D, S817D* single mutant.. This suggests that most of the deficiency is due to a decrease in initial capture, rather than in reversal.

The extension of D-loops stabilizes the nascent three-strand intermediates, and it is possible that defects in capture may also be related to defects in D-loop extension. D-loop extension from the newly paired 3’ end can be measured using the same strains (**Figure 3**D and Extended View **Figure 8**AB). As with the D-loop capture assay, there was only a modest difference from WT in the ability of *rad54S816A, S817A* to extend D-loops, and there was a significant loss (20-fold) of D-loop extension in the *rad54S816D, S817D* strain at 6 hours (**Figure 3E and Extended View Figure 8**CDE). This is compared to a 100-fold loss in the *rad54* strain. Surprisingly, in the *rad54S816D, S817D sgs1* strain, there was now a 100-fold loss in D-loop extension, suggesting that while D-loop capture was slightly improved, there was insufficient extension to reach the second restriction site (**Figure 3E**). Importantly, this means that the *rad54S816D, S817D* strains are competent for D-loop extension if the D-loop is initially stabilized. The primary defect in this mutant occurs at the D-loop capture stage.

The extended D-loops can also be characterized by their ssDNA and dsDNA content (**Figure 3F**). Because this is a break-induced replication system, the amount of dsDNA produced reflects the amount of lagging-strand synthesis. This value is low in WT but is 3-4-fold higher in *sgs1* strains (19). We also observed this increase in dsDNA in a *sgs1* strain (**Figure 3G**). Surprisingly, this increase was lost in the *rad54S816A, S817A sgs1* strain (**Figure 3G**). This result suggests a genetic interaction between *RAD54* and *SGS1* during the repair synthesis phase, and a slight defect in the *rad54S816A, S817A* mutant. It should be noted that the extension values for *rad54* and *rad54S816D, S817D sgs1* were so low that this type of analysis could not yield reliable results.

### Recombination between alleles

The *rad54S816D, S817D* mutant had diminished D-loop capture and failed to promote survival during ectopic recombination. This type of recombination is generally less efficient than recombination between sister chromatids or homologous chromosomes and, therefore, may be more sensitive to subtle changes in recombination efficiency (86). Consequently, we investigated the impact of the *rad54S816A, S817A,* and *rad54S816D, S817D* alleles on recombination between homologous chromosomes. We used a reporter assay based on the *ADE2* gene in diploid *S. cerevisiae* (90–93). Each copy of chromosome XV has a different inactive allele of *ade2*; one copy contains an *I-Sce1* nuclease site (*ade2-I*), and the other has an inactivating mutation located within the gene (*ade2-n*) (**Figure 4A**). Upon induction of the nuclease and formation of a double-strand break, both sister chromatids are cut, and repair occurs with the *ade2-n* allele located on the homologous chromosomes. Types of gene conversion events can be determined by reconstitution of the *ADE2* genes. If long tract repair occurs, then the gene conversion event will acquire the deleterious mutation from the homolog, and the yeast colony will appear red. If short tract repair occurs, the mutation will not be acquired, and a functional *ADE2* gene will be reconstituted, resulting in a white colony (**Figure 4B**). Each sister can be repaired via a long tract or short tract gene conversion event, and due to independent assortment, a colony can be red (two long tracts), white (two short tracts), or sectored (one long tract and one short tract).

In our hands, sectored colonies are the most common outcome in WT, occurring approximately 65-70% of the time. This was also observed in the *rad54S816A, S817A* mutant (**Figure 4C**). However, with the *rad54S816D, S817D* allele, sectored colonies accounted for only about 20-25% of the population, with most colonies appearing solid red. Approximately 95% of these colonies were recombinants rather than uncut colonies. This observation is based on re-induction with galactose. Recombinants should be able to survive on galactose plates since the DSB is repaired and the original cut site is lost after repair. We also observed a 3-fold increase in the percentage of solid white colonies, though the significance of this finding is unclear. We quantified the ratio of long-tract to short-tract conversion by counting individual conversion events. For example, a solid red or white colony counts as two events, while a sectored colony counts as one short-tract and one long-tract gene conversion. According to this analysis, the WT results in a long-to-short tract ratio of 1.7 (**Figure 4D**). A similar ratio was calculated for the *rad54S816, S817A* mutants (**Figure 4D**), while the ratio was two times higher (3.4) in the *rad54S816D, S817D* mutant. From these data, we conclude that the mutant form of Rad54 results in more frequent long-tract gene conversion events.

The helicase Sgs1 is known to shorten gene conversion tracts. From our previous experiments, we identified a novel genetic interaction between Rad54 and Sgs1. Therefore, we generated *SGS1* deletion strains with *RAD54*, *rad54S816A,S817A*, and *rad54S816D,S817D*. Deletion of *SGS1* alone resulted in a minor increase in gene conversion tract length (**Figure 4**CD). However, *rad54S816A, S817A sgs1* strains now had a 2-fold increase in the number of long tract conversions, which was comparable to the *rad54S816D, S817D* strain (**Figure 4**CD). In the *rad54S816D, S817D sgs1* strain, there was a further increase in long tract gene conversion to 4 to 5-fold more than WT (**Figure 4**CD). This data further supports a synthetic interaction between Rad54 and Sgs1, and the increase in long tract gene conversion is likely due to reduced stability of the primary strand invasion intermediates.

The reporter strains used in these experiments also carry antibiotic markers downstream of the *ade2* genes. The segregation of these markers can be used to determine whether the recombination results in a Crossover (CO), Non-Crossover (NCO), or Break-induced Replication (BIR) outcome (**Figure 4**AB). Outcomes were analyzed from sectored (**Figure 4E**) and solid red colonies (**Figure 4F**). The same outcomes were also pooled to examine the total recombination outcomes (**Figure 4G**). For the sectored colonies, there was a reduction in CO outcomes for both the *rad54S816A, S817A* and *rad54S816D, S817D* alleles (**Figure 4E**). This occurred due to an increase in NCO outcomes in the *rad54S816A, S817A* strain, and an increase in BIR in the *rad54S816D, S817D* strain. The most dramatic impact was observed in solid red colonies, where the *rad54S816D, S817D* allele primarily repaired through NCO and BIR outcomes (**Figure 4F**). Upon analyzing the total recombination outcomes, we observed a general trend of increased NCO and BIR outcomes (**Figure 4G**). These outcomes are stronger in the *rad54S816D, S817D* strain, but also occur in the *rad54S816A, S817A* strain.

Upon deletion of *SGS1* in the *rad54S816A, S817A* strains, there was an increase in NCO outcomes as well as BIR. This came at the expense of CO outcomes (**Figure 4**EFG). This phenotype is enhanced in the *rad54S816D, S817D* strain, resulting in a near-complete loss of CO outcomes (**Figure 4**EFG). A substantial increase in BIR or BIR-like outcomes balances this loss. The increase in long-tract gene conversion, the increase in BIR, and the loss of CO outcomes likely reflect a reduction in second DNA-end engagement. According to this interpretation, the data suggest that stabilization of the primary strand invasion intermediate by Rad54 and Sgs1 may be a necessary step in properly capturing the second end of DNA for double Holliday junction formation.

### Rad54 S816DS817D is defective in binding dsDNA

Through our genetic analysis, we determined that substituting Aspartic acid for S816 and S817 disrupts D-loop stabilization *in vivo* and may also disrupt second-end DNA capture. To further characterize the defects in protein activity, we expressed and purified Rad54 S816A/S817A and Rad54 S816/DS817D and tested their biochemical properties.

Rad54 is a dsDNA-dependent ATPase, and we measured ATP hydrolysis efficiency for WT, Rad54 S816A/S817A, and Rad54 S816D/S817D. There was no difference between WT and Rad54 S816A/S817A (**Extended View Figure 9A**). However, the Rad54S816D/S817D protein poorly hydrolyzed ATP (**Extended View Figure 9A**). This loss of activity could be due to an inactive enzyme or reduced dsDNA binding. Rad51 further stimulates Rad54 ATPase activity by 3-5-fold. To determine if the disruption of ATP hydrolysis was due to a dead enzyme or binding to dsDNA, we added Rad51 to the ATPase reaction. In the WT and Rad54 S816AS817A, there was the expected stimulation of hydrolysis. Unexpectedly, the addition of Rad51 to the Rad54 S816DS817D mutant resulted in a 20-fold stimulation of activity, making hydrolysis levels comparable to the WT (**Extended View Figure 9A**). These data suggest that the Rad54 S816DS817D mutant may have difficulty binding to dsDNA in the absence of Rad51, but this is resolved when Rad51 is present.

Because the Rad54 S816D/S817D mutant shows reduced ATPase activity, we next tested its ability to bind dsDNA using an electromobility shift assay. We measured a 2.5-fold reduction in binding to dsDNA for the Rad54 S816D/S817D mutant compared to the WT (K_d_ 27 nM versus 68 nM) (**Extended View Figure 9**BC). A small reduction in the apparent K_d_ for Rad54 S816A/S817A was also observed (K_d_ 27 nM versus 36 nM) (**Extended View Figure 9**BC). Rad54 is known to form an oligomer (94). A reduction in oligomer formation could cause an apparent defect in binding to dsDNA. The cooperativity of protein binding to DNA would be reflected in this. Therefore, we measured the hill coefficient from our binding plots. We observed no difference in binding cooperativity (h constant 3.7 versus 3.6 of WT and Rad54 S816D/S817D, respectively), suggesting that the defect is due to direct binding to dsDNA rather than oligomerization.

To more quantitatively evaluate the defect in binding, we used single-molecule DNA curtains (95,96) to monitor the stability of GFP-Rad54, GFP-Rad54 S816A/S817A, and GFP-Rad54 S816D/S817D (**Figure 5**AB). These experiments measure the dissociation rate of the Rad54 protein, the velocity at which it moves along dsDNA, and the distance it travels (**Figure 5**AB). To accurately determine the rate of Rad54 dissociation, we first validated that the loss of protein signal was due to dissociation rather than photobleaching. We measured the photobleaching rate by monitoring GFP loss when the laser was left on without shuttering. The loss of GFP-Rad54 signal was then measured with shuttering of the laser. If the rate of signal loss exceeds the rate of photobleaching with shuttering, we are measuring dissociation events. Rad54, Rad54 S816AS817A, and Rad54 S816DS817D all dissociated faster than photobleaching (**Figure 5C**). These measurements were converted to dissociation rates, and it was determined that Rad54 S816D/S817D dissociated more rapidly than WT or Rad54 S816A/S817A (**Figure 5D**). This is consistent with reduced dsDNA binding by the Rad54 S816D/S817D mutant.

Rad54 dissociation was measured with the buffer flow on, which applies a force to the proteins and can affect the rate and direction of movement along the dsDNA. Rad54 can translocate in either direction or can switch direction as it moves along dsDNA (29). The direction depends on the DNA strand used as the tracking strand. In our experiments, when Rad54 translocates toward the barriers, it does so against the flow of buffer or a resisting force. In contrast, if movement is toward the pedestals, then an assisting force is present. We were surprised to observe that there was no difference in the velocity of Rad54 movement with or against the force of flow between the WT, Rad54 S816AS817A, and Rad54 S816DS817D (**Figure 5E**), suggesting no general defect in translocation by the Rad54 S816D, S817D mutant.

There were differences in the observed distance translocated both with and against the flow (**Figure 5F**). This difference was small between the WT and Rad54 S816A/S817A (4759 ± 2591 bp versus 5671 ± 3391 bp when an assisting force was applied, and 8930 ± 4136 bp versus 6954 ± 3744 bp when a resisting force was applied). However, a much larger difference was observed between the WT and Rad54 S816D S817D (4759 +/-2591 bp versus 7820 +/-4859 bp with an assisting force, 8940 +/-4136 bp versus 5740 +/-3667 bp when with a resisting force). There was also a difference in the direction of movement along the DNA. Surprisingly, 75 to 80% of WT molecules translocated against the buffer flow (**Figure 5G**). This may be due to the DNA in that direction being more tense, which could promote directional movement (97). In contrast, only 15-20% of Rad54 S816D/S817D molecules translocated against the flow (**Figure 5G**). These observations are consistent with weaker binding by Rad54 S816D/S817D to the DNA. Lower affinity allows the force of buffer flow to push Rad54 S816D/S817D 1 dimensionally along the DNA, resulting in longer tracks only in the direction of flow, but maintaining similar velocities.

### PSCs with Rad54 S816D817D are less active than WT

The Rad51-ssDNA-Rad54 complex is referred to as the presynaptic filament (PSC) (**Figure 6A**). We previously used PSC reconstitution in combination with DNA curtains to monitor Rad54 activity in the context of the PSC, and this complex is considered active during homology search and strand exchange in HR. The combination of Rad51 and Rad54 in the PSC can exert greater force on the DNA than Rad54 alone (97) and may make it more resistant to buffer flow. Therefore, we tested the activity of PSCs reconstituted with Rad54, Rad54 S816A/S817A, and Rad54 S816D/S817D (**Figure 6B**).

In WT and the Rad54 mutants, 80-85% of PSCs moved against the flow and exhibited considerably higher velocities and distances during translocation against the buffer flow (**Figure 6**CDE). These observations are all consistent with increased affinity for the DNA relative to Rad54 alone. In contrast to Rad54 alone, there was a difference in the translocation velocity between PSCs with Rad54, Rad54 S816AS817A, and Rad54 S816DS817D. The velocity of Rad54 S816AS817A was 85% of WT, and Rad54 S816DS817D was 60% of WT. These differences were observed only when the PSCs moved against the flow, and no differences were observed when they moved with the flow (**Figure 6**CDE). These differences are consistent with weakened binding by PSCs carrying mutant Rad54. We did not observe differences in the distance of translocation in either direction, suggesting that all PSCs are more stable than Rad54 alone. We used a previously established technique to monitor the number of Rad54 molecules bound in each PSC (29,92). Under conditions where flow is constant, there is no difference in the number of Rad54 molecules bound per PSC. This suggests that the reduction of velocity against flow may occur through different mechanisms. From these experiments, we conclude there is only a minor difference in the movement of the PSC with Rad54 S816D/S817D.

We reasoned that defects in PSC movement could result from a loss of force generation by Rad54 on the donor DNA. The application of force to the donor DNA by the PSC results in the DNA’s deformation. This distortion can turn the donor into a substrate for the ssDNA binding protein RPA (29,96) (**Figure 7A**). Therefore, a reduction-in-force application should result in differences in RPA binding within the PSC. For these experiments, the PSC was formed and allowed to function with the flow off for 5 minutes before the flow was restored, and the RPA signal was monitored. Approximately 63% (53/83) of WT PSCs recruited RPA. In contrast, only 34% (47/138) of Rad54 S816D/S817D PSCs had RPA associated (**Figure 7B**). In the Rad54 S816A/S817A mutant, there was a slight reduction in RPA, as well (48%, 58/122). This is consistent with the differences in activity observed for the PSC with Rad54 S816A/S817A, which also translated to the physiological setting during allelic recombination and repair synthesis.

These results indicate that mutation of this region of Rad54 reduces DNA remodeling, a consequence of the force applied to the DNA. Rad54 exerts force on the donor DNA by establishing multiple points of contact (97), and this activity is enhanced when DNA tension is reduced. Therefore, we reasoned that turning off the buffer flow would allow PSCs with WT Rad54 to accumulate more Rad54. We measured the estimated number of Rad54 molecules per PSC. This number was 2-fold higher than in our flow experiments with WT and Rad54 S816A/S817A. Notably, the number of Rad54 S816D/S817D molecules was significantly lower (**Figure 7C**). The change in Rad54 molecules bound in the mutant is likely due to the failure to isolate local donor DNA. It is unlikely to be a self-recruitment issue, given the equivalent cooperativity observed by EMSA. Therefore, we conclude that rad54 S816D/S817D has reduced capacity to remodel local donor DNA structures, likely due to decreased ability to overcome pre-existing force on the DNA.

Previously, we hypothesized that applying force to DNA may promote the stabilization of nascent D-loops. The *rad54S816D, S817D* mutant was defective in D-loop formation *both in vivo* and *in vitro*, particularly under force application to dsDNA. We directly measured whether the Rad54 S816D/S817D mutant was defective in forming D-loops *in vitro* using a previously established assay (93,96). The Rad54 S816AS817A mutant was able to promote D-loop formation while the Rad54 S816DS/817D protein did not (**Extended View Figure 9D**). This observation is consistent with the lack of D-loop formation observed *in vivo.* It supports a model in which the application of force to DNA is essential for stabilizing nascent D-loops.

## Discussion

In this study, we identified a universally conserved interaction in Rad54, the mutation of which leads to the separation of distinct Rad54 functions. Disruption in contact between the two RecA lobes of Rad54 results in a functional protein that fails to form D-loops both *in vivo* and *in vitro*. However, it remains active in removing Rad51 that is pathologically bound to dsDNA and can complement *rad54* strains in the presence of certain DNA-damaging agents. Our data indicate that the Rad54 mutant is defective in pump activity and has severely diminished capacity to catalyze D-loop formation.

### A Pump or a Motor

Brownian ratchets are molecular machines that harness thermal fluctuations in proteins to perform useful directional work (98). This is usually achieved by binding a cofactor, such as ATP, which traps specific protein conformations. ATP-dependent ratchets can be categorized as motors or pumps. Motors use ATP hydrolysis to translocate along a linear or rotational track, such as a DNA helix. At the same time, pumps move molecules to create energy gradients. In both cases, motor slippage can reduce movement efficiency or disrupt the concentration gradient. The definition of Rad54 as a motor is obvious, and ATP hydrolysis allows physical movement along the DNA. However, Rad54 can also promote DNA underwinding by isolating short DNA segments (97). These isolated regions store energy in the form of underwound DNA, and by this definition, Rad54 can be considered a pump.

Our data suggest that the Rad54 S816D/S817D mutation has little impact on linear translocation along DNA and instead affects Rad54’s pump function. This occurs by disrupting the force applied to the DNA. To act as a pump and create a superhelical density gradient in the DNA, Rad54 must form multiple contacts with the substrate (97). If a single point of contact is lost in this organization, then the gradient will dissipate. The instability of the Rad54 S816D/S817D mutant could cause it to dissociate from dsDNA more frequently, leading to a loss of topological isolation.

A second possibility is that the mutation generates a slippery ratchet, allowing leakage of isolated turns in the underwound region of DNA. Slippage could occur during Rad51-mediated strand exchange. Previous work has demonstrated that Rad51 invasion results in the addition of negative turns to the DNA (99,100). After enough turns are added to the DNA, the DNA collapses into a plectoneme or melts to relieve torsional stress. The inclusion of Rad54 in the PSC promotes DNA backbone melting as turns are added (97). Slippage of Rad54 would likely reduce the efficiency of this step-in strand exchange (**Figure 7D**). This step precedes the removal of Rad51 from the newly formed joint molecule, thus identifying a novel intermediate. Therefore, several of the activities assigned to Rad54 during HR are likely to depend on balancing its motor, removing Rad51 from dsDNA, and on pump functions that isolate underwound regions of dsDNA.

### Specific mechanism of Rad54 separation of function mutant

Based on structural modeling, the substitution of S816 and S817 to Aspartic acid likely disrupts a contact that forms between the two RecA lobes of Rad54. Disruption affects Rad54’s affinity for dsDNA. Similar structures and protein sequences are identified in all Rad54/Rdh54 (Rad54B) family members, as well as the closely related replication fork remodeling enzyme ATRX (101). The interaction is not found in other members of the Snf2 family of chromatin remodelers. A second function of Rad54 is to suppress the binding of Rad51 to dsDNA, stemming from the ability of Rad54 to remove Rad51 from dsDNA. This is generally considered a pathogenic state characterized by Rad51 overexpression, a condition that occurs in many types of human cancers (38). In our hands, the *rad54S816D, S817D* alleles suppressed phenotypes associated with pathogenic RAD51 overexpression, indicating that these mutants are separation-of-function. The ability to remove Rad51 from dsDNA has been proposed as the mechanism by which D-loops form. This model explains how Rad51 is removed from DNA, enabling DNA extension and repair (47,49).

The Rad51 removal model is widely accepted but has been challenged by models that propose Rad54 alters topology as a mechanism for D-loop formation. By identifying this separation-of-function mutant, we can clarify these two models. While the *Rad54S816D, S817D* mutant has a severe defect in D-loop capture, it still promotes DNA extension at a level consistent with the amount of initial capture. No further defects are observed at extension, because this version of the protein is competent to remove Rad51. This observation is supported by the loss of extension in the *rad54S816D, S817D sgs1* double mutant, suggesting that the small amount of extension that occurs in the *rad54S816D, S817D* is lost. Stabilization is regulated by topology, and extension is regulated by Rad51 removal. Therefore, both activities are required in sequence to explain Rad54’s activity at D-loop structures and the transition to DNA synthesis.

Previous phosphoproteomic experiments identified Rad54 S816 as a kinase site. We were unable to identify the specific kinase or context in which this residue might be phosphorylated. However, given that a phosphomimetic mutation at this site results in a separation-of-function mutant, it is possible that phosphorylation is cell-cycle stage-dependent and effectively balances the need to promote strand exchange with the need to remove Rad51 from dsDNA. This mechanism has been proposed for the kinase NEK1 in human cells (102), and a kinase that phosphorylates Rad54 at S816 may collaborate with NEK1 to regulate Rad54 function. Based on our analysis, the most likely kinase would come from the Polo-like kinase family. However, further research is needed to validate this hypothesis.

### The long and short of it

Mutations in Rad54 resulted in generally higher levels of long-tract gene conversion than WT during allelic recombination. The phenotypes appear distinct from those associated with previous Rad54 hypomorphic alleles, which primarily affect translocation (96). These mutants affected recombination outcomes but did not alter gene conversion tracts. The length of gene conversion tracts is a function of DNA polymerase activity (103). The *ade2-n* mutation is located ∼950 bp upstream of the *I-Sce1* site in the *ade2-I* gene. Therefore, a higher proportion of strand exchange outcomes will need to occur on the 5’ side of this mutation. Exchange would require longer ssDNA resection products or longer Rad51 filaments to invade upstream of the *ade2* mutation. These events could occur due to reduced D-loop capture efficiency, requiring longer Rad51 filaments, or a higher density of short Rad51 filaments on ssDNA to overcome the limitations imposed by reduced Rad54 function. Alternatively, the increase in long tract repair could be due to high levels of short extension events, followed by disruption and reinvasion (23). This is consistent with the increase in BIR and NCO outcomes observed in our experiments, and with the reduction in DNA extension observed *in rad54S816D, S817D sgs1* strains.

The high incidence of long tract repair, accompanied by increased NCO and BIR outcomes, may also reflect increased replication. In our system, full BIR outcomes require replication to the end of the chromosome. Shorter BIR events may become indistinguishable from NCO outcomes generated by SDSA and would appear as non-crossover outcomes. A current hypothesis is that Srs2 prevents the formation of toxic intermediates that may arise from ssDNA generated during BIR (22). The strand separation activity of Srs2 is not required during HR, and disruption of this activity is not synthetic lethal with *RAD54* (65). However, removal of Rad51 from the ssDNA generated during BIR prevents the accumulation of toxic intermediates. These outcomes have the potential to be more deleterious and involve significant opportunity for non-allelic genomic rearrangements. Therefore, the remodeling of the local DNA environment by Rad54 and Sgs1 may serve a protective function during HR by preventing potentially toxic repair intermediates from forming during the BIR pathway.

The loss of CO outcomes and increase in BIR are symptomatic of reduced second-end engagement. Moreover, the near-complete loss of CO outcomes suggests that the double Holliday junction (dHJ) repair pathway may be lost, as this is the most likely repair pathway leading to CO formation. This severe outcome only occurred in the *rad54S816D, S817D sgs1* strain, but a general trend of CO loss and BIR increase is observed in the *sgs1*, *rad54S816A, S817A sgs1*, and *rad54S816D, S817D sgs1* strains. A significant reduction in DNA extension, like that observed in the *rad54S816D, S817D sgs1* strain, will also diminish second-end engagement, potentially impacting second-end capture (104). Both helicases may contribute to the stable formation of this critical intermediate. Biochemically, the reduction of Rad54 binding to dsDNA and the topological isolation of donor DNA regions may contribute to the stable formation of the dHJ. It is less clear how Sgs1 may contribute to this outcome mechanistically. dHJ formation in the absence of Rad54 is observed in many organisms, and during meiosis (105). In these scenarios, alternative proteins may remodel DNA in a manner like Rad54, thereby stabilizing primary-strand exchange intermediates to promote second-end capture and Holliday junction formation. Therefore, the underlying cause of increased second-end engagement is likely regulation of DNA topology.

An alternative possibility is that the inability to regulate topology at strand-exchange intermediates leads to reduced resistance to anti-crossover factors, such as Sgs1-Top3-Rmi1 (STR) (86,106). Rad54 and STR have both been implicated in branch migration during HR. These outcomes are consistent with observations in the *rad54 S816A, S817A,* and *rad54S816D, S817D* single mutants. The act of branch migration generates significant topological stress along the DNA, with positive supercoils forming ahead of the branch. Therefore, branch migration to promote dissolution might be inhibited by Rad54’s ability to isolate a DNA domain topologically. Alternatively, the lack of Rad54 binding and stabilization could reduce cleavage by the Mus81-Mms4 complex, thereby facilitating the formation of crossovers and half-crossovers (22,91). These possibilities remain speculative, and future work will be needed to understand the impact of Rad54-mediated DNA remodeling on recombination outcomes.

## Conclusions

In this study, we identified a universally conserved mutation in Rad54 that reduces binding to dsDNA. This caused a severe defect in D-loop formation *in vivo* but did not alter the protein’s ability to remove Rad51 from double-stranded DNA. From this, we hypothesize that the separation-of-function mutant fails to stabilize initial D-loop formation but remains competent to remove Rad51 from the newly formed D-loop. This apparent separation of function enables us to better understand Rad54’s sequential action at initial strand-exchange intermediates, thereby unifying two existing models in the field. Future work should focus on understanding how Rad54’s loss of DNA-binding affects outcomes in human cell lines and at DNA replication forks.

## Data availability

All data is available upon request.

## Acknowledgements

We would like to acknowledge Eric Alani and Eric C. Greene for critical reading of the manuscript. We want to thank Wolf Heyer, Lorraine Symington, and Jim Haber for providing the parent reporter strains used in this study. We would also like to thank the members of the Cornell R3 group and the Crickard laboratory for their helpful input during the project’s development.

## Author contributions

JH performed the initial screen, performed experiments, analyzed single molecule experiments, and helped write the manuscript. DM performed single molecule experiments, helped analyze the single molecule experiments, and provided input on writing. AX helped with the initial screen and performed imaging of cellular foci. LP performed spot assays for Rad54 mutants. JBC performed allelic recombination experiments, guided the research, provided funding, and wrote the manuscript with input from all authors. This work is supported by NIGMS R35142457 and American Cancer Research Scholar Grant RSG-25-1410641-01-DMC to JBC.

## Materials and Methods

### Yeast strains construction

Strains for the initial spot assay were BY4741 and generated by centromeric plasmid expression in the *rad54* strain or by gene replacement, as indicated in the context. Strains for the ectopic recombination assay were TGI354 (86) and were generated by gene replacement or centromeric plasmid expression in the *rad54* strain, as indicated in the context. Strains for the *in vivo* D-loop capture (DLC) and extension (DLE) assay were generated in the WDHY5511(19,88) background by gene replacement. Strains for tetrad dissection were derived from the W303 background and generated by gene replacement. Strains for red/white allelic recombination were generated by gene replacement from parent strains LSY2205 and LSY2202 (91). Strains for the allelic recombination assay were in W303, TGI354, and WDHY5511 backgrounds, which were also used in the spot assay as indicated.

### Protein purification

Rad54, Rad54 S816AS817A, Rad54 S816DS817D, Rad51, and RPA-mCherry were purified as previously described (29). In brief, a protease-deficient yeast strain was transformed with GFP-GST-Rad54, GFP-GST-Rad54 S816AS817A, or GFP-GST-Rad54 S816DS817D on a 2 µ plasmid under the control of the Gal1/10 promoter. Cells were grown in YNB (–Ura) plus 3% glycerol and 2% lactic acid. When the cells reached an OD_600_ of 1.5, expression was induced by adding 2% galactose for 6 hours. Cells were harvested by centrifugation and stored at –80°C.

Cell pellets were resuspended in Rad54 resuspension buffer (30 mM Tris–HCl [pH 7.5], 1 M NaCl, 1 mM EDTA, 10% glycerol, 10 mM BME (β-mercaptoethanol), protease inhibitor cocktail and 2 mM PMSF). Cells were disrupted by manual bead beating, and the lysate was clarified by centrifugation at 26,500 x g for 1 hour. The lysate was fractionated by ammonium sulfate (AS) precipitation. AS was gradually added with mixing to a final concentration of 20% followed by centrifugation at 10,000 x g for 10 minutes. The supernatant was discarded, and the AS concentration was raised to 50% followed by centrifugation at 10,000 x g for 10 min. The protein pellet was resuspended in PBS (phosphate buffered saline) plus 1M NaCl and 10 mM BME. The resulting re-suspended protein was then bound to pre-equilibrated GST resin in batch for 1 hour at 4°C. The GST resin was washed twice with PBS plus 1000 mM NaCl, and twice with PBS plus 500 mM NaCl. The protein was eluted in 20 mM glutathione in PBS plus 500 mM NaCl. The peak fractions were pooled and then applied to a Sephacryl S–300 High Resolution gel filtration column pre-equilibrated with Rad54 SEC buffer (30 mM Tris–HCl [pH 7.5], 500 mM NaCl, 1 mM EDTA, 10% glycerol, and 10 mM BME). The peak was pooled and dialyzed against Rad54 SEC buffer plus 50% glycerol and stored at –80°C in single-use aliquots.

6xHis–SUMO–Rad51 was transformed into *Escherichia coli* BL21 (DE3) Rosetta2 cells and grown to an OD_600_ of 0.4 to 0.6 at 37°C. Expression was induced by addition of 0.5 mM IPTG for 3 hours at 37°C. Cells were harvested and stored at –80°C. Cells were lysed by freeze–thaw in Cell Lysis Buffer (CLB: 30 mM Tris–HCl [pH 8.0], 1 M NaCl, 10% glycerol, 10 mM imidazole, 5 mM BME, and protease inhibitor cocktail). Crude lysates were sonicated for 6 pulses of 30 seconds on and 2 minutes off, then clarified by centrifugation at 26,500 x g. The extract was precipitated with 50% AS and centrifuged at 26,500 x g for 10 minutes. The pellet was resuspended in CLB and bound to 1 mL of pre–equilibrated Ni–NTA resin for 1 hour with rotation at 4 °C. The resin was washed three times with CLB and eluted in CLB plus 200 mM imidazole. The protein was mixed with 400 units of the SUMO protease Ulp1 and dialyzed overnight at 4°C into Rad51 buffer (30 mM Tris–HCl [pH 8.0], 150 mM NaCl, 1 mM EDTA, 10% Glycerol, 10 mM imidazole). The 6xHis–SUMO tag and SUMO protease were removed by passing the dialyzed proteins over a second 1 mL Ni–NTA column. The purified Rad51 was then stored at –80°C in single-use aliquots.

### Multiple Sequence Alignment

Rad54-like eukaryotic sequences were retrieved by BLASTP (NCBI, nr database; organism filter: Eukaryota) using *S.c.*Rad54 as the query. Redundant entries and obvious fragments were removed.

Sequences were aligned with Clustal Omega, the alignment was trimmed with trimAl, and sequence logos were generated from the trimmed alignment using the Python package Logomaker.

### Yeast spot growth assay

*RAD54* and its designed mutants were expressed using a centromere vector, pRS415, and transformed into the BY4741 *rad54* strain or other background strains as indicated, followed by selection on YNB (–Leu) + 2% dextrose plates. Transformed cells were grown overnight in YNB (– Leu) + 2% dextrose medium. The following day, the overnight cultures were diluted to an OD_600_ of 0.3 and grown to an OD_600_ of 1.0. Cells were then serially diluted and spotted on YNB (-Leu) + 2% dextrose plates containing either no drug, 0.01% methyl methanesulfonate (MMS), or 0.02% MMS. Other drugs were also used, including 0.02 µg/mL 4-NQO, 0.04 µg/mL 4-NQO, 10 µM camptothecin (CPT), and 20 µM CPT as indicated. Plates were incubated at 30 °C for 3 days and imaged at 72 hours. BY4741 strains with RAD54 mutants were also generated via gene replacement. The procedures for the spot assay for these strains were the same as for the centromeric-expressed strains, except that the overnight culture was grown and diluted in Yeast extract + Peptone (YP) + 2% dextrose medium. The serially diluted cells were spotted on YP + 2% dextrose containing either no drug or the drugs mentioned above. Plates were incubated at 30 °C for 2 days and imaged at 48 hours. For *RAD51* or *rad51I345T* on a pYES plasmid, cells were transformed. Cultures were grown overnight in YNB (-Ura) + 2% dextrose and diluted the next day. After cells reached an OD_600_ of 1.0, they were serially diluted onto YNB (-Ura) +2% dextrose or YNB (-Ura) +2% galactose and incubated at 30 °C for 48 hours. They were then images.

### Ectopic recombination assay

The wildtype strain used for this study was described in (86). The genotypes for modifying these strains are listed in Extended Table 1. A single colony was picked and grown in YP + 3% glycerol + 2% lactate medium to log phase. The culture was diluted and plated on YP + 2% dextrose and YP + 2% galactose plates, respectively. Plates were incubated at 30 °C for 2 to 3 days, and the number of colonies was counted. The viability rate was calculated by dividing the colony number on the YP + 2% galactose plate by that on the YP + 2% dextrose plate. The mean and standard deviation were calculated for multiple independent experiments as indicated. For centromeric expression of RAD54 and its mutants in *rad54* strains, a single colony was picked and grown in YNB (-Leu) + 2% dextrose overnight. On the second day, the culture was diluted 10-fold into YNB (-Leu) containing 3% glycerol and 2% lactate and grown at 30 °C for 6 hours. Then the culture was diluted and plated onto YNB (-Leu) + 2% dextrose and YNB (-Leu) + 2% galactose, respectively. After being incubated at 30 °C for 3 to 4 days, the numbers of colonies were determined, and the viability was calculated by dividing the colony number on the galactose plate by that on the dextrose plate.

### Allelic Recombination assay

The assay was performed by growing the appropriate strain overnight in YP + 2% raffinose. The next day, cells were diluted to an OD_600_ of 0.2 and allowed to reach an OD_600_ of 0.4-0.5, then *I-Sce1* expression was induced by adding 2% galactose. Cells were allowed to grow for an additional 1.5 hours, then plated onto YP + 2% dextrose (YPD) plates and grown for 48 hours. After 48 hours, they were placed at 4 °C overnight to enable further development of the red color. The number of white, red, and sectored colonies was then counted, followed by replica plating onto YPD + hygromycin B (200 µg/ml) and YPD + nourseothricin (67 µg/ml, clonNat) for analysis of recombination outcomes. Strains were also replica plated on YNB (-Ura/-Met) + 2% dextrose to ensure proper chromosome segregation and YNB (-adenine sulfate) +2% galactose to assay for recombinants versus uncut DNA. The data were analyzed by counting sectored colonies and assessing colony survival across different antibiotic sensitivities. The data for each category was then divided by the total population of sectored colonies. The standard deviation between biological replicates was analyzed for at least six independent experiments from different crosses.

### Yeast cell imaging

To immobilize yeast cells, an agarose pad was used. A 1% agarose solution was prepared by adding agarose in S media (YNB plus 0.5% ammonium sulfate, then autoclaved) and microwaving to dissolve. An amino acid mix was added to a final concentration of 1x to ensure yeast cells could grow on the agarose pad. The dissolved agarose was kept warm on a 65 °C metal heater. One clean microscope slide was taped at both ends, and 30 µL of 1% agarose was applied to the center of the slide. Another clean and untapped microscope slide was immediately placed on top, and gentle pressure was applied to ensure the agarose spread evenly. Then the slide sandwich was placed on an ice bag for approximately 30 seconds to allow the agarose to solidify. The top slide was carefully slid out, leaving the agarose pad intact on the bottom slide. The agarose pad was allowed to dry for 3 minutes before 3 µL of properly diluted yeast cells was added. A coverslip was then placed over the cells. A Nikon Eclipse Ti microscope equipped with a 488-nm laser. A Plan APO 60XA/1.20 WI objective and an ANDOR Zyla-5.5-CL3 camera were used to observe and capture yeast cells.

### Tetrad dissection

Strains used for tetrad dissection were generated by gene replacement in the W303 background and are listed in Extended Table 1. *MATa* strains and *MATα* strains were first mixed on YPD plates and incubated for 4 hours at 30 °C, then re-streaked on YNB (-Leu) + 2% dextrose + 0.25 g/L G418 plates to select for both *TRP1* and *KanMX* markers. Diploid cells were re-streaked onto sporulation plates (1% KOAc, 0.1% yeast extract, 0.05% dextrose, 2% Agar) and grown for 2 days at 30 °C. One loopful of sporulated cells was digested in zymolyase at 37 °C for 9.5 minutes before tetrad dissection. The 10x zymolyase solution for tetrad dissection was prepared by dissolving 25 mg zymolyase in 500 μL of spheroplasting buffer (recipe for spheroplasting buffer described in the DLC assay section). For a working tetrad dissection solution, 2.5 μL of 10x zymolyase solution was diluted into 25 μL sterile water. The digestion was quenched by adding 250 μL sterile water and placing the cells on ice for at least 5 minutes. For dissection, 10 μL of digested spores were added onto one side of a YPD plate, which was then placed vertically to allow the drop flow across the plate. Plates were stored at 4 °C for 2 days before tetrad dissection. Dissection was performed on a dissecting microscope. One tetrad was separated into four spores, which were then placed individually on the plate with ∼5 mm spacing. The spores were grown at 30 °C for 2 days before survival rates were assessed.

### ATPase assay

A commercially available ADP-GLO kit (Cat No. V6930, Promega) was used to measure ATP hydrolysis activity. ATP hydrolysis reaction was performed in HR buffer (20 mM Tris-OAc [pH 7.5], 50 mM NaCl, 10 mM Mg(OAc)_2_, 200 ng/μl BSA, 1 mM DTT, and 10% Glycerol) and contained 1 mg/ml sheared salmon sperm DNA, 20 nM Rad54, and 200 nM Rad51 (if added).

### Electrophoresis mobility shift assay (EMSA) for Rad54

An Atto647N-labeled 90-mer oligo was annealed with an unlabeled complementary oligo to form a labeled 90-bp dsDNA substrate. The oligo sequences are available in Extended Table 3. The binding reaction was performed in EMSA buffer (35 mM Tris-Cl [pH 7.5], 3 mM MgCl_2_, 50 mM KCl, 1 mM DTT, 10% glycerol). The final DNA concentration was 10 nM, and proteins were titrated to be 0, 6.25, 12.5, 25, 50, 100, 150, and 200 nM as final concentrations for Rad54 wildtype and Rad54 S816DS817D. Rad54 S816AS817A was titrated to be 0, 6.25, 12.5, 25, 50, 100, and 139 nM as final concentrations. The DNA and proteins were incubated at 30 °C for 5 min and then resolved by 8% Native-PAGE in 0.5x TBE buffer (44.6 mM Tris, 44.5 mM boric acid, 1 mM EDTA, 8% acrylamide / bis-acrylamide (37.5:1), 0.1% APS, 0.1% TEMED) running in 0.5x TBE buffer (44.6 mM Tris, 44.5 mM boric acid, 1 mM EDTA). The hill curve of Rad54-DNA binding was fitted through neutcurve (107).

#### *In vitro* D-loop assay

D-loop formation experiments were performed in HR buffer (30 mM Tris-OAc [pH 7.5], 50 mM NaCl, 10 mM MgOAc_2_, 1 mM DTT, 0.2 mg/ml BSA) using an Atto647N-labeled DNA duplex (15 nM) with homology to the pUC19 plasmid. Rad51 (300 nM) was incubated with recipient DNA at 30 °C for 15 minutes. The resulting Rad51 filaments were mixed with indicated concentrations of Rad54, RPA (500 nM), and pUC19 plasmid (0.3 nM). Reactions were quenched at the indicated time points and treated with 1 unit of Proteinase K at 37°C for 20 minutes. The reactions were then resolved by electrophoresis on a 0.9% agarose gel and imaged for fluorescence using a Typhoon imager.

### Flow cell construction

Metallic chrome patterns were deposited on quartz microscope slides with predrilled holes for microfluidic line attachment by electron beam lithography to generate flow cells. After metal deposition, a channel was created by placing a small piece of paper between the drill holes and covering the two-sided tape. The paper was excised to make the flow chamber, and a glass coverslip was fixed to the tape. The chamber was sealed by heating to 165 °C in a vacuum oven at 25 mmHg for 60 min. Flow cells were then completed by hot-gluing IDEX nano ports over the drill holes on the opposite side of the microscope slide from the coverslip.

### Single molecule experiments

All single molecule experiments were conducted on a custom-built prism-based total internal reflection microscope (Nikon) equipped with a 488-nm laser (Coherent Sapphire, 100 mW), a 561- nm laser (Coherent Sapphire, 100 mW), a 640-nm laser (Coherent Obis, 100 mW) and two Andor iXon EMCCD cameras. DNA substrates for DNA curtains experiments were made by attaching a biotinylated oligo to one end of the 50 kb Lambda phage genome and an oligo with a digoxigenin moiety on the other. This enabled double tethering of the DNA between the chrome barriers and the chrome pedestals, as previously described. As for single tethered DNA, if specified in the context, flow cells were attached to a microfluidic system, and sample delivery was controlled using a syringe pump (KD Scientific). Three-color imaging was achieved by two XION 512 ×512 back-thinned Andor EM-CCD cameras and alternative illumination using a 488 nm laser, a 561 nm laser, and a 640 nm laser at 25% power output. The lasers were shuttered, resulting in a 200- msec delay between each frame. Images were collected with a 200-msec integration time. Translocation velocity and distances were measured in HR Buffer. Channel bleed-through is prevented by shuttering of the laser lines, emission filters, and the use of complementary fluorophores. In this case GFP is not activated by the 561 or 647 laser lines, and the 488 or 561 laser lines do not activate Atto647N. mCherry labelled the 488-laser line can poorly activate fluorophores. However, the emitted light is split by a dichroic mirror and filtered through a band- pass and long-pass filter to block wavelengths of light above a certain cutoff. This is sufficient to prevent mCherry signal bleed through into the 488 channel.

### Analysis of dsDNA translocation

The velocity and track length for GFP-Rad54 molecules were measured by importing raw TIFF images as image stacks into ImageJ. Kymographs were generated by pointing around a fluorophore signal and defining the whole individual dsDNA molecules where the fluorophore bound to a region of interest (ROI) using a home-made script. Data analysis was performed from the kymographs. The start of translocation was defined when the GFP-Rad54 molecule moved > 2 pixels. Pauses were defined as momentary stalls in translocation that lasted 2–4 frames. Termination was defined by molecules that did not move for > 10 frames. Velocities were calculated using the following formula [(Y_f_ –Y_i_) ×1031.96 bp / [|X_f_ –X_i_|]) ×frame rate]; where Y_i_ and Y_f_ correspond to the initial and final pixel position and X_i_ and X_f_ correspond to the start and stop time (in seconds). Negative values stand for translocations moving from barrier to pedestals, which was assisted by a flow. Positive values stand for translocations against flow directions.

### Dissociation rate measurement

The dissociation rate of DNA-GFP-Rad54 was measured by initiating data collection, followed by the injection of 10nM GFP-Rad54 in HR buffer onto single-tethered DNA curtains to allow proteins to bind to DNA. The flow rate was maintained at 0.2 mL/min to allow for initial binding and prevent further binding events. Images were collected with a 100 msec integration time at 200 msec intervals for a duration of 5 minutes for the bleaching experiment, or at 1 sec intervals for a period of 15 minutes for the dissociation experiment. The fluorescent signal decay in both experiments was fit to an exponential function, A = A_max_*exp(-k*t). The difference between the k values obtained from bleaching experiment and dissociation experiment was used to calculate the dissociation rate: k_off_ = k_dissociation_ - k_bleaching_.

### DLC assay

DLC assay was performed as described before (19,88) . In brief, yeast cells were grown in a 5 mL YP + 2% dextrose + 4% adenine sulfate medium overnight. The second day, the culture was diluted by 10-fold in 5 mL YP + 3% glycerol + 2% lactate + 4% adenine sulfate and grown for around 8 hours. Then the culture was inoculated into 100 mL of YP + 3% glycerol + 2% lactate + 4% adenine sulfate medium with an initial OD_600_ ≈ 0.006 and grown for 16 hours. A 5x psoralen stock solution (0.5 mg/mL trioxsalen in 200-proof ethanol) was made in a 50-mL aluminum foil-covered falcon tube and dissolved on a shaker at room temperature overnight with gentle rocking. Next day, the culture should have an OD_600_ at 0.3-0.8. 7.5 OD_600_ of cells was collected as time 0 control, centrifuged at 2,246 x g, 4 °C for 5 minutes.

The cell pellets were resuspended in 1x psoralen buffer. The 1x psoralen buffer was prepared by diluting 5x psoralen in 200-proof ethanol before collecting cells. The resuspended cells were plated in a 60 mm x 15 mm petri dish, put 2-3 cm below a UV light source with the lip removed atop a pre-chilled metal block. The cell samples were exposed under the UV light for 10 minutes with gentle shaking to crosslink DNA. The cells were transferred to a 15-mL Falcon tube. The petri dish was rinsed with TE1 solution (50 mM Tris-Cl [pH 8.0], 50 mM EDTA), and the TE1 buffer was poured over the cells. The cells were then centrifuged at 2,246 x g, 4 °C for 5 minutes again. The supernatant was properly disposed of, and the pellets were stored at -20 °C. Galactose was added to the culture to a final concentration of 2% to induce DSBs. Cells were collected at the designed time points as described above. For lysis, the cell pellets were thawed on ice, then resuspended in spheroplasting buffer (0.4 M sorbitol, 0.4 M KCl, 40 mM sodium phosphate buffer [pH 7.2], 0.5 mM MgCl_2_) and transferred to a 1.7 mL microfuge tube.

The cells were spheroplasted in zymolyase solution (2% glucose, 50 mM Tris-Cl [pH 7.5], 5 mg/mL zymolyase 100T) at 30 °C for 20 minutes. The cells were washed with spheroplasting buffer three times at 2,500 x g and restriction enzyme buffer (RE buffer: 50 mM potassium acetate, 20 mM Tris-acetate, 10 mM magnesium acetate, 1 mg/mL BSA) at 16,000 x g three times. The pellets were resuspended in 1.4x RE buffer, either alone or with a hybridization oligo to restore the *EcoR*I restriction sites, and stored at -80 °C. The DNA was solubilized by incubating the cells with 0.1% SDS at 65 °C for 13 minutes. 1% Triton X-100 quenched the SDS. The DNA was digested by 20 U *EcoR*I at 37 °C for 1 hour. The restriction enzyme was deactivated by incubating the DNA with 1.5% SDS at 55 °C for 10 minutes. The cells were returned to ice, and SDS was quenched by the addition of 6% Triton X-100. Ligation buffer (50 mM Tris-HCl [pH 8.0], 10 mM MgCl2, 10 mM DTT, 2.5 μg/mL BSA, 1 mM ATP, pH 8.0, 8 U T4 DNA ligase) was added to perform the ligation reaction at 16 °C for 1 hour and 30 minutes. 25 μg/mL protease K was added to digest the enzymes at 65 °C for 30 minutes. DNA was extracted by adding phenol:chloroform:isoamyl alcohol and vortexing. The upper water phase was moved and incubated with a tenth volume of sodium acetate and a volume of isopropanol at room temperature for 30 minutes and centrifuged at 21,130 x g, 4 °C for 10 minutes to get DNA precipitation. The DNA pellets were dried at 37 °C and dissolved by incubating with 1x TE buffer (10 mM Tris-Cl [pH 8.0], 1 mM EDTA) at 37 °C for 1 hour. The DNA was used as a qPCR template with the primers listed in Extended Table 3. DLC chimera content was calculated by [DLC amplification efficiency]^[-Cp_(DLC)_], and the intramolecular ligation product content was calculated by [intramolecular ligation amplification efficiency]^[-Cp_(ligation)_]. The final DLC signal was calculated as DLC chimera content divided by intramolecular ligation product content.

### DLE assay

DLE assay was performed as described before(19,88). The procedures were similar with DLC assay except for (1) 2.5 OD_600_ of cells were collect at each time point; (2) DNA crosslinking was omitted and cell pellets were washed by TE1 buffer twice at 2,246 x g; (3) hybridization oligos were different and listed in Extended Table 4; (4) DNA was solubilized by 1% SDS at 65 °C for 15 minutes; (4) DNA digestion was performed by adding 20 U HindIII; (5) qPCR was performed using some different oligos listed in Extended Table 5.

