## Supplemental Figures for "Rad54 separation of function mutation highlights unique roles during homologous recombination"

### Extended View

**Extended Table 1**

Strains used in the study

| Strains | Genotype | Source or reference |
|---|---|---|
| BY4741 (WT) | | Dharmacon |
| BY4741-*rad54*Δ | *rad54*Δ | Dharmacon |
| BY4741 | *RAD54::rad54S816A-KANMX* | This study |
| BY4741 | *RAD54::rad54S817A-KANMX* | This study |
| BY4741 | *RAD54::rad54S816A,S817A-KANMX* | This study |
| BY4741 | *RAD54::rad54S816D-KANMX* | This study |
| BY4741 | *RAD54::rad54S817D-KANMX* | This study |
| BY4741 | *RAD54::rad54S816D,S817D-KANMX* | This study |
| WDHY5511 | *ura3::A-HOcs, lys2::A, trp1::GAL-HO-hphMX, his3D200, can1-100, leu2–3, 112, ade2–1, RAD5* | Piazza et al 2020 |
| JBC423 (WDHY5511) | *rad54::KANMX* | This study |
| JBC281 (WDHY5511) | *RAD54::RAD54-KANMX* | This study |
| JBC367 (WDHY5511) | *RAD54::rad54S816A,S817A-KANMX* | This study |
| JBC369 (WDHY5511) | *RAD54::rad54S816D,S817D-KANMX* | This study |
| JBC380 (WDHY5511) | *sgs1::HISIII* | This study |
| JBC388 (WDHY5511) | *RAD54::rad54S816A,S817A-KANMX, sgs1::HISIII* | This study |
| JBC478 (WDHY5511) | *RAD54::rad54S816D,S817D-KANMX, sgs1::HISIII* | This Study |
| TGI354 | *TGI 354 JKM 146 (arg5,6::MATa::HPH)* | Ira et al 2003 |
| JBC425 (TGI354) | *rad54::KANMX* | This study |
| JBC357 (TGI354) | *RAD54::RAD54-KANMX* | This study |
| JBC359 (TGI354) | *RAD54::rad54S816A,S817A-KANMX* | This study |
| JBC361 (TGI354) | *RAD54::rad54S816D,S817D-KANMX* | This study |
| Protease deficient yeast strain | *MATα leu2 trp1 ura3–52 prb1–1122 his3::pGAL1–GAL* | Crickard et al 2020 |
| 11C (LYS2202) | *MATα ade2-I lys2:GAL-ISCEI his3:HphMX4* | Mazon et al. 2010 |
| 15D (LYS2205) | *MATa ade2-n his3:NatMX4 met22:klURA3* | Mazon et al. 2010 |
| JBC106 | *11C-RAD54::RAD54-KanMX* | Keymakh et al 2022 |
| JBC105 | *15D-RAD54::RAD54-KanMX* | Keymakh et al 2022 |
| JBC413 | *11C-RAD54::rad54S816AS817A-KanMX* | This Study |

| | | |
|---|---|---|
| JBC414 | *15D-RAD54::rad54S816AS817A-KanMX* | This Study |
| JBC479 | *11C-RAD54::rad54S816DS817D-KanMX* | This Study |
| JBC417 | *15D-RAD54::rad54S816DS817D-KanMX* | This Study |
| JBC 434 | *11C-RAD54::RAD54-KanMX sgs1::HIS3* | This Study |
| JBC 436 | *15D-RAD54::RAD54-KanMX sg1::HIS3* | This Study |
| JBC 430 | *11C-RAD54::rad54S816AS817A-KanMX sgs1::HIS3* | This Study |
| JBC 432 | *15D-RAD54::rad54S816AS817A-KanMX sgs1::HIS3* | This Study |
| JBC522 | *11C-RAD54::rad54S816DS817D-KanMX sgs1::HIS3* | This Study |
| JBC444 | *15D-RAD54::rad54S816DS817D-KanMX sgs1::HIS3* | This Study |
| JBC501 | *W303 MATa srs2::TRP1* | This Study |
| JBC506 | *W303 MATα RAD54::rad54S816D,S817D-3xHA-KanMX* | This Study |
| JBC510 | *W303 MATa RAD54::RAD54-3xHA-KanMX* | This Study |
| JBC514 | *W303 MATa rad54::KanMX* | This Study |
| JBC490(WDHY 5511) | *rad51I345T-LEU2* | |
| JBC517 (WDHY5511) | *Rad54 S816D, S817D-KanMx rad51I345T-LEU2* | |
| **Diploids** | | |
| JBC106xJBC105 | | |
| JBC413xJBC414 | | This Study |
| JBC479xJBC417 | | This Study |
| JBC434xJBC436 | | This Study |
| JBC430X JBC432 | | This Study |
| JBC444xJBC522 | | This Study |

**Extended Table 2**
**Plasmid in the study**

| Backbone | Construction | Source |
|---|---|---|
| pRS305 | *RAD54-KANMX* | This study |
| pRS305 | *rad54S816A-KANMX* | This study |
| pRS305 | *rad54S816D-KANMX* | This study |
| pRS305 | *rad54S816AS817A-KANMX* | This study |
| pRS305 | *rad54S816DS817D-KANMX* | This study |

| | | |
|---|---|---|
| pRS415 | *RAD54* | Crickard et al. 2020 |
| pRS415 | *rad54S816A* | This study |
| pRS415 | *rad54S816D* | This study |
| pRS415 | *rad54S816AS817A* | This study |
| pRS415 | *rad54S816DS817D* | This study |
| pRS415 | *rad54T85A* | This study |
| pRS415 | *rad54T85E* | This study |
| pRS415 | *rad5T132A* | This study |
| pRS415 | *rad5T132E* | This study |
| pRS415 | *rad5T155A* | This study |
| pRS415 | *rad5T155E* | This study |
| pRS415 | *rad5S214A* | This study |
| pRS415 | *rad5S214D* | This study |
| pRS415 | *rad5T231A* | This study |
| pRS415 | *rad5T231E* | This study |
| pRS415 | *rad5S303A* | This study |
| pRS415 | *rad5S303D* | This study |
| pRS415 | *rad54S318A* | This study |
| pRS415 | *rad54S318D* | This study |
| pYES | *GST-GFP-RAD54* | Crickard et al, *Cell*, 2020 |
| pYES | *GST-GFP-rad54S816,AS817A* | This study |
| pYES | *GST-GFP-rad54S816,DS817D* | This study |

**Extended Table 3**
Oligos used in the study

| Name | Sequence | Purpose |
|---|---|---|
| 90-mer DNA | Atto647N-GATGTTCTGCTGGATATGCACTTTTCCGGGCTGACGTACACCGTGCTCAGCCTGTTTTCAGCGATCCGGATATGCATCCGCTGGATTTC | Single Molecule imaging |
| olWDH1760 | CAGCGGGCTTGCAGAAGTTG | To amplify genomic DNA at *ARG4* |
| olWDH1761 | GGCCAATTAGTTCACCAAGACG | To amplify genomic DNA at *ARG4* |
| olWDH1766 | GTTTCAGCTTTCCGCAACAG | To quantify DSB induction |
| olWDH1767 | GGCGAGGTATTGGATAGTTCC | To quantify DSB induction |
| olWDH2009 | CACCACTTTGCCATTCAACAC | To amplify at the downstream site of elongation |
| olWDH2010 | TGCTCGGAGATTACCGAATC | To amplify at upstream site of the |

| | | |
|---|---|---|
| | | DSB, used with olWDH2009 to quantify DLE signal |
| olWDH2011 | TGCGAGGTTTTCTTGGTCAG | Used with olWDH2009 to quantify *Hin*dIII recognition site restoration |
| olWDH2012 | CGAGGCATATTATGGTGAAGG | Used with olWDH2010 to measures *Hin*dIII recognition site restoration |
| olWDH2052 | ATGTGCCTTCCTACCGCTC | To quantify intramolecular ligation efficiency of *Hin*dIII-derived fragments |
| olWDH2053 | TCAAGCGTGGTTACATTCCTTAC | To quantify intramolecular ligation efficiency of *Hin*dIII-derived fragments |
| olWDH2007 | TCTGCTCGGAGATTACCGAATCAAAAAAATT TCAAAGAAACCGGAATCAAAAAAAAGAAC AAAAAAAAAAAAGATGAATTGAAAAGCTTT ATGGACCGAC | To restore *Hin*dIII site |
| olWDH2046 | AATCTTTGTGAAGCTTCGCAAGTATTCATTTT AGACCCATGGTGGAACCCTAGTGTTGAATG GCAAAGTGGTGATAGAGTTCATAGAATTGGT CAGTAT | To restore *Hin*dIII site |
| olWDH1762 | ACTTCGAATTTCGGCACTTC | To quantify intramolecular ligation efficiency of *Eco*RI-derived fragments |
| olWDH1763 | CGATGAAACGTTAAGTGACCAC | To quantify intramolecular ligation efficiency of *Eco*RI-derived fragments |
| olWDH1764 | AGAGCGGTCAGTAGCAATCC | To amplify at the upstream of DSB |
| olWDH1765 | CACACGCGAAAAACCGCC | To amplify at the upstream of the donor DNA, used with olWDH1764 to quantify DLC signal |
| olWDH2019 | CTTTAACCGGACGCTCGA | To quantify psoralen crosslinking efficiency |

| | | |
|---|---|---|
| olWDH2020 | TTGAGTTTATTGCTGCCGTC | To quantify psoralen crosslinking efficiency |
| olWDH1768 | AGGAGCACAGACTTAGATTGG | Used with olWDH1764 to measure *Eco*RI recognition site restoration |
| olWDH1770 | CGAAATCATCTTCGGTTAAATCCAAAACGGCAGAAGCCTGAATGAAACATATGAACCAATTGGAGGACGTCAATGAATTCTGGGGATCCATTGCATTTTT | To restore *Eco*RI site |

### Extended View Figure 1:

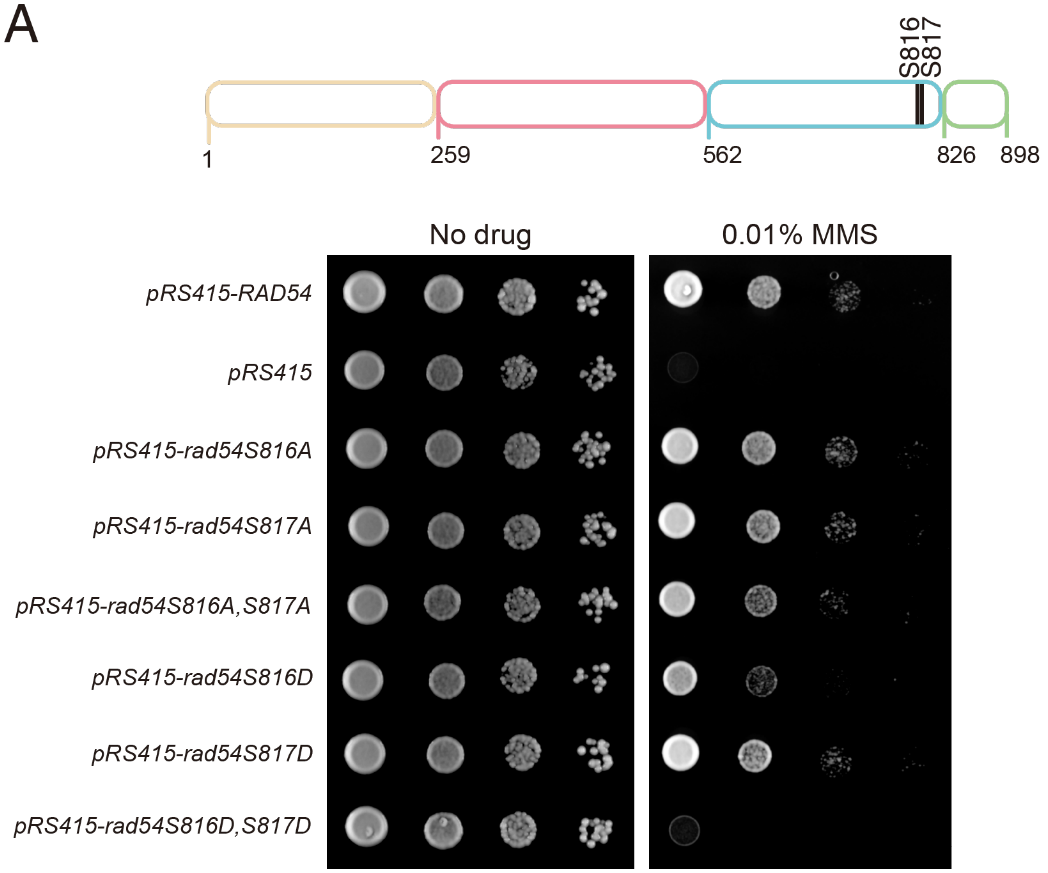


**Extended View Figure 1: *rad54S816DS817D* is more deficient than *rad54S816D***
**(A).** Serial dilution spot assay to determine the sensitivity of yeast strains with single amino acid substitutions at Rad54 S816 and S817 or double mutations at the same positions. Serine was mutated to either Alanine or Aspartic acid.

### Extended View Figure 2:

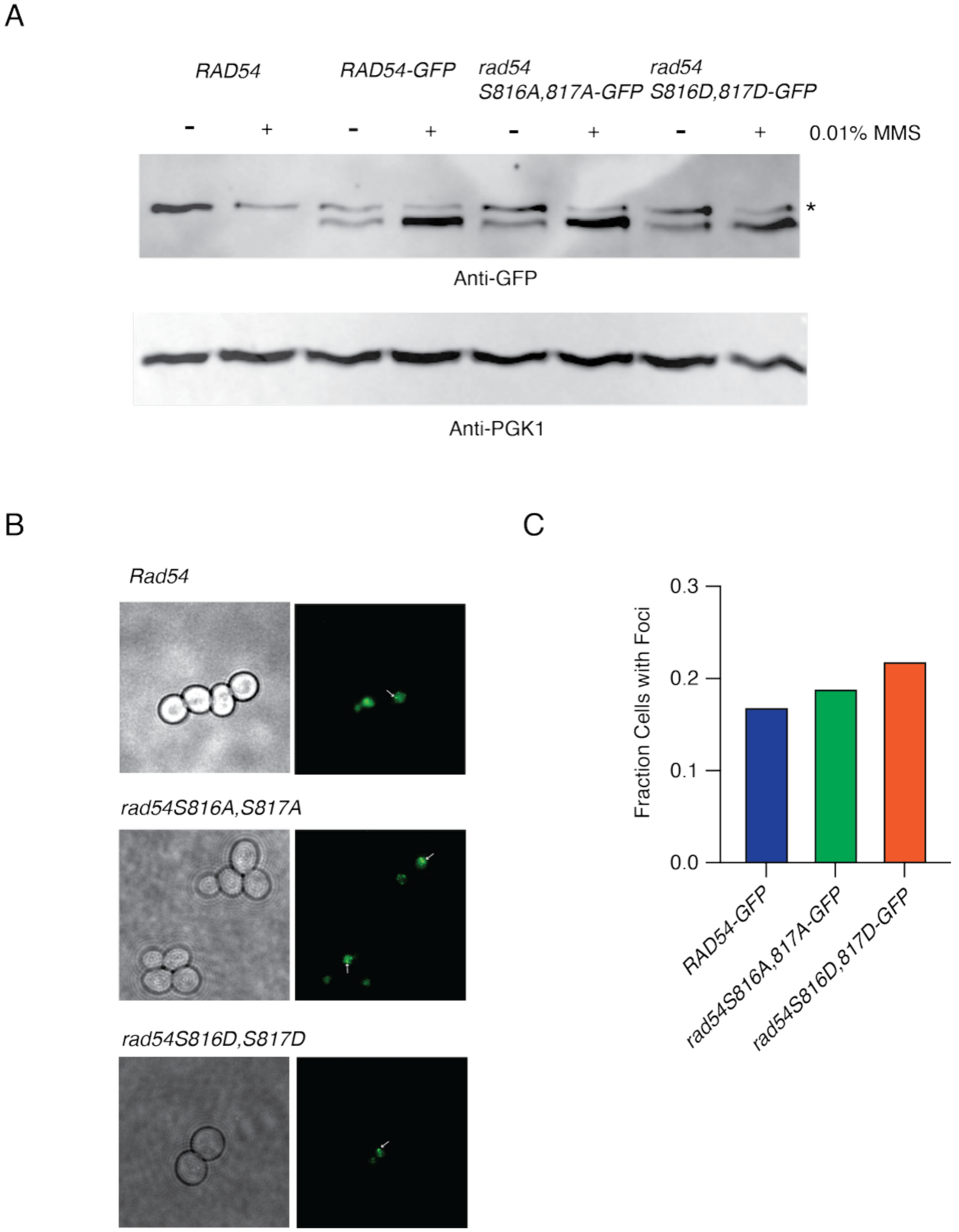


**Extended View Figure 2: Mutant forms of Rad54 are expressed in cells**
**(A).** Representative western blot illustrating the expression of *RAD54-GFP*, *rad54S816A, S817A-GFP*, and *rad54S816D, S817D-GFP* with and without 0.01% MMS. PGK1 is used as a loading control. The Asterix represents a non-specific band**(B).** Fluorescent microscope images illustrating that *RAD54-GFP*, *rad54S816A, S817A-GFP*, and *rad54S816D, S817D-GFP* form foci in response to MMS treatment. **(C).** Bar graph quantifying the percentage of cells that form foci in response to MMS treatment.

#### Extended View Figure 3

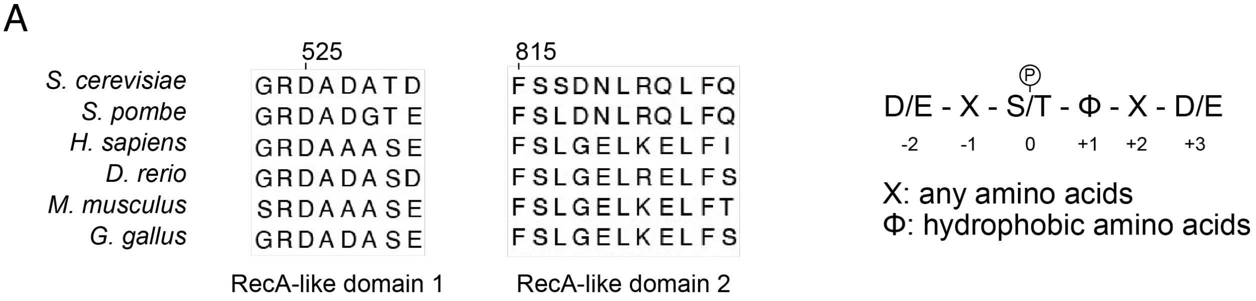


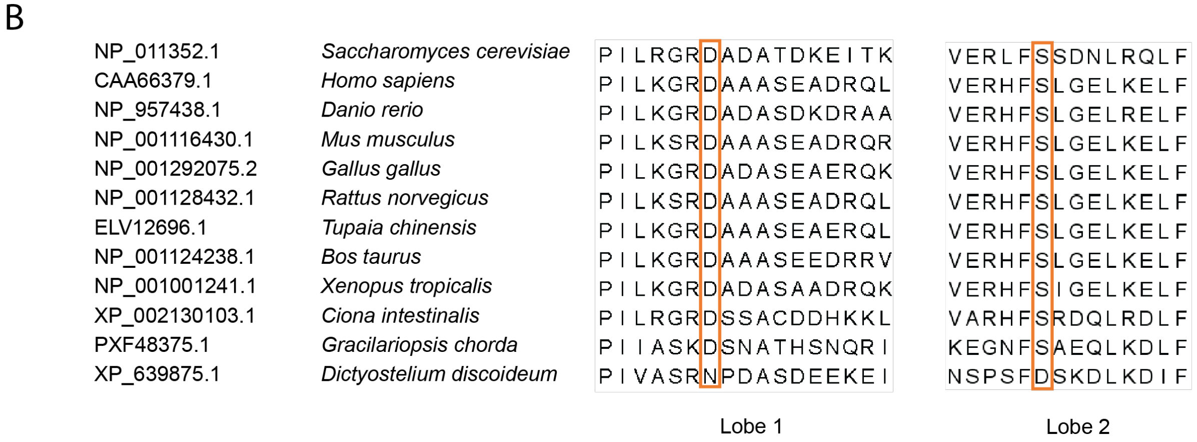


**Extended View Figure 3: Excerpt from multiple sequence alignment**

**(A).** Multiple sequence alignment of Rad54 interacting region (Left) and consensus kinase sequence for Polo-like-kinase (Right). The human RAD54 matches the consensus sequence.

**(B).** An excerpt from the Rad54 multiple sequence alignment. Shown are the two interacting regions on lobes 1 and 2, respectively. Included in this alignment in *Dictyostelium discoideum,* which has a naturally occurring Aspartic acid in place of Serine on lobe 2.

### Extended View Figure 4

A

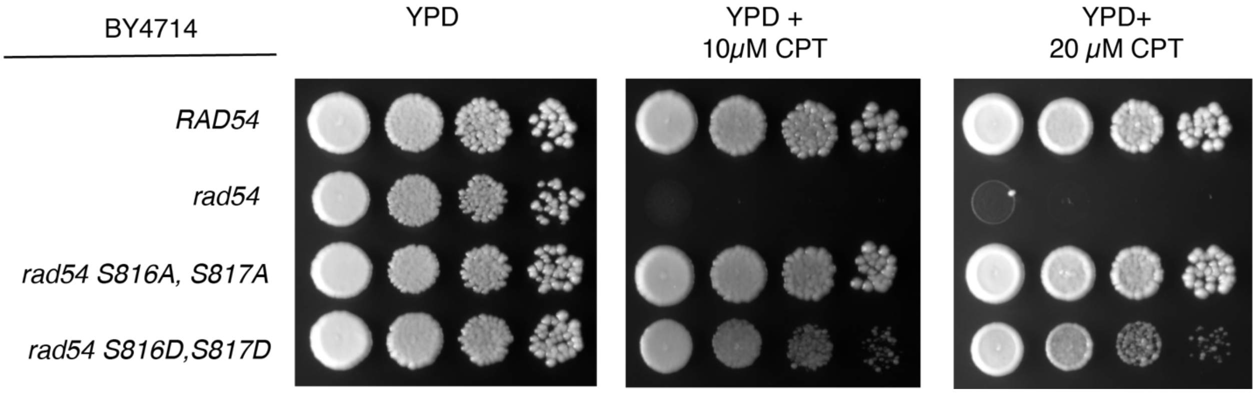


**Extended View Figure 4: BY4741 are less sensitive to CPT in *rad54S816D,S817D***
**(A).** Spot assay illustrating *RAD54*, *rad54*, *rad54S816A, S817A,* and *rad54S816D, S817D* sensitivity to CPT.

### Extended View Figure 5

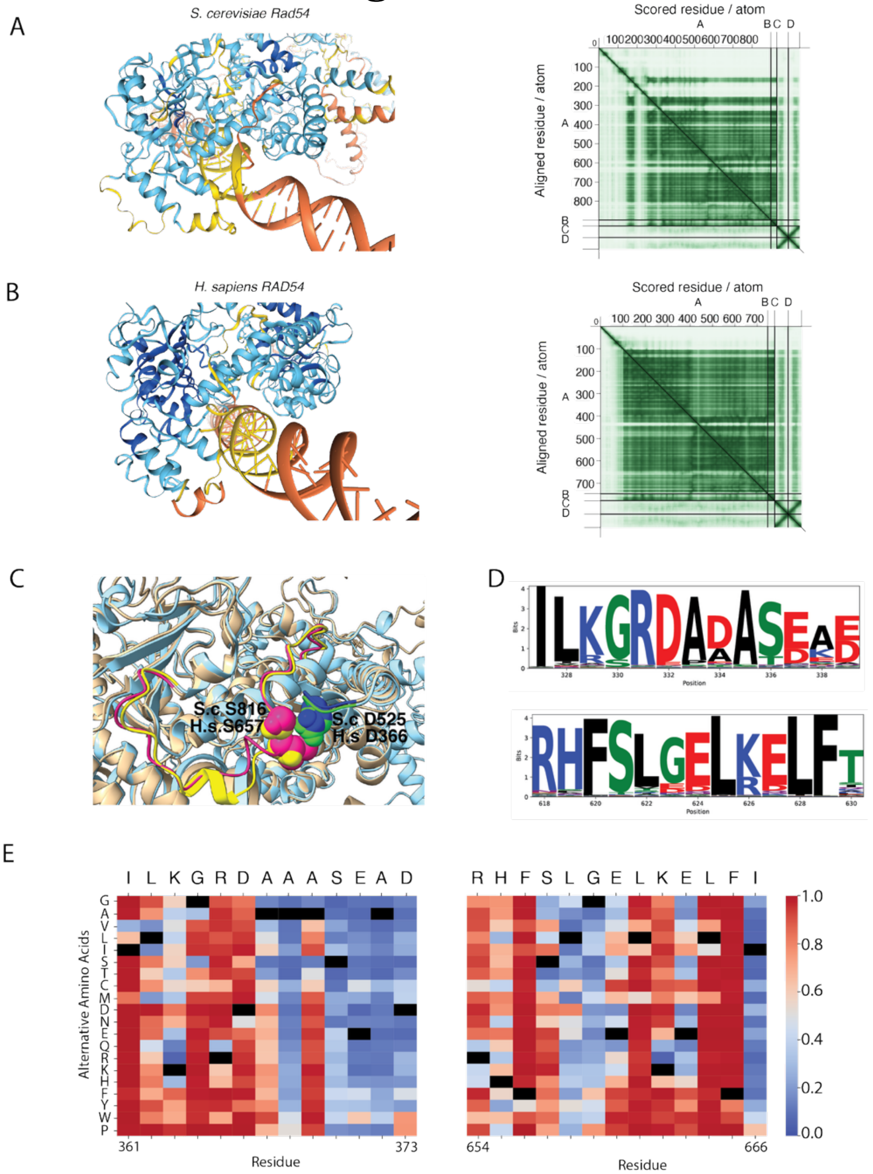


**Extended View Figure 5: Computational analysis of potential phosphorylation sites on Rad54**

**(A).** AlphaFold3 model for *S. cerevisiae* Rad54 bound to dsDNA, colored for confidence with the pLDDT formation (Left). PAE plot for *S. cerevisiae* Rad54 AlphaFold structure (Right). **(B).** AlphaFold3 model for *H. sapiens* RAD54 bound to dsDNA, colored for confidence with the pLDDT formation (Left). PAE plot for *H. sapiens* RAD54 AlphaFold structure (Right). **(C).** Structural superposition of *S. cerevisiae* Rad54 and *H. Sapiens* RAD54. The predicted interacting residues are shown as space-filling spheres. **(D).** Protein sequence logos generated from 1224 eukaryotic Rad54 sequences, illustrating the sequence conservation in these two regions using logo maker. **(E).** Sequence view of interacting region of *H. sapiens* RAD54 using a predicted pathogenicity plot for individual amino acid substitutions generated via AlphaFold.

### Extended View Figure 6

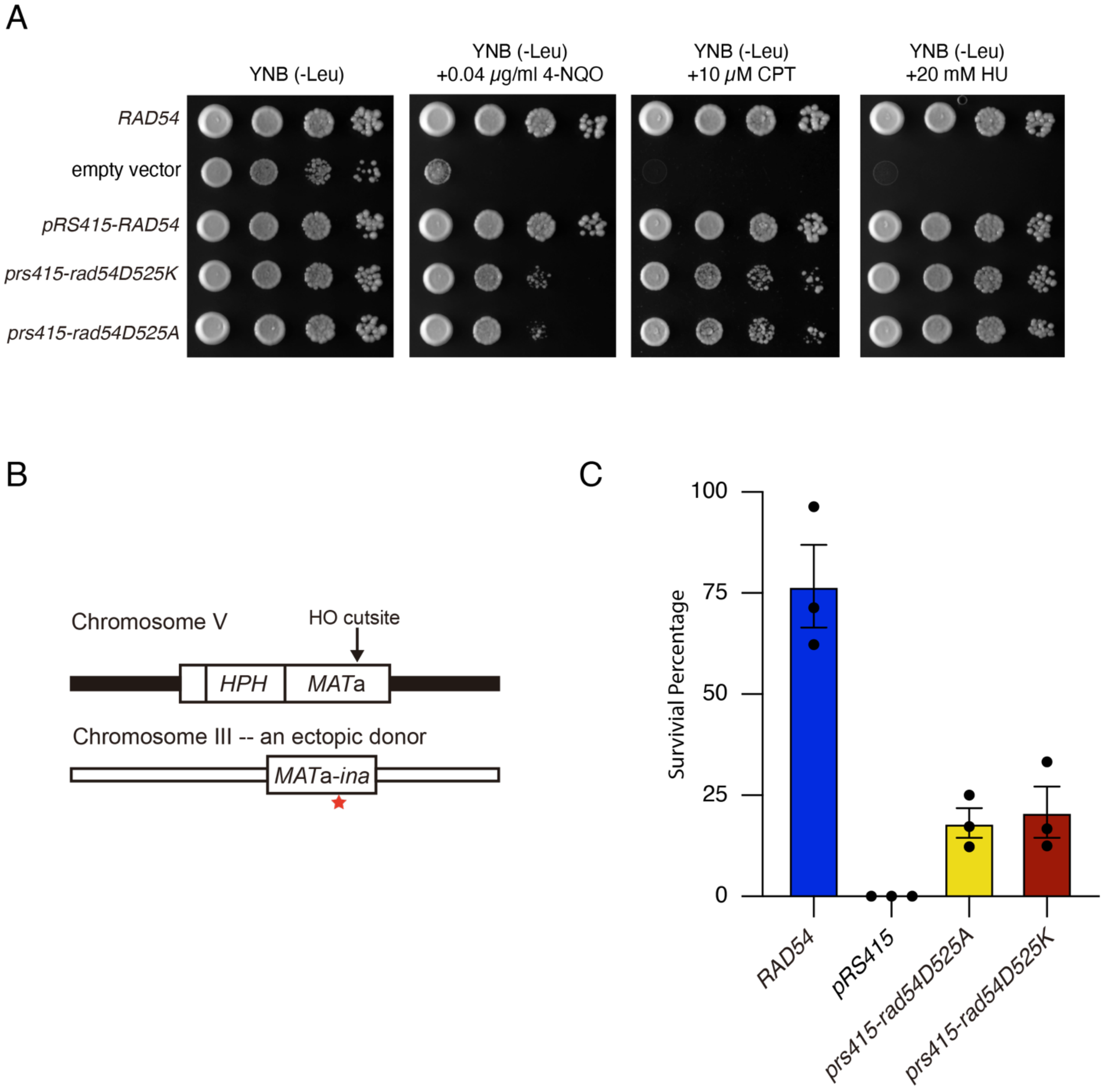


**Extended View Figure 6:** *rad54D525A* and *rad54D525K* phenocopy phosphomimic mutants **(A)** Yeast spot assays testing the ability of *rad54D525A* and *rad54D525K* to complement 4-NQO, CPT, and HU sensitivity. **(B).** Schematic diagram for ectopic double-strand break repair assay. **(C).** Bar graph representing the survival percentage of DSB-induced cells in TGI354 *rad54* expressing *pRS415-RAD54*, empty vector, *pRS415-rad54D525A*, and *pRS415-rad54D525K*. The bars represent the mean, and the error bars represent the standard error measurement from three independent experiments.

### Extended View Figure 7

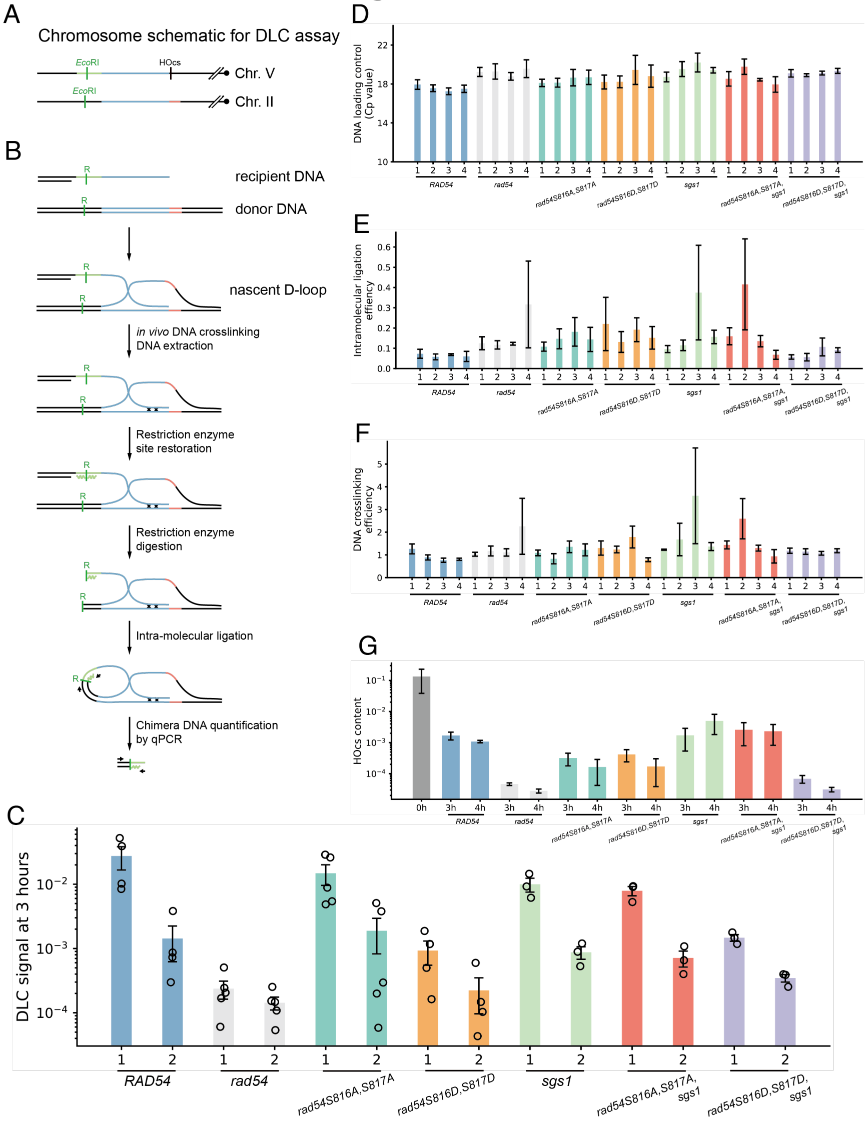


**Extended View Figure 7: Supplementary information for the DLC assay**

**(A).** Chromatin schematic for the DLC assay. **(B).** Rationale for the DLC assay. **(C).** DLC signal at 3 hours. Column set 1 shows results with a hybrid oligo added; column set 2 shows results without adding a hybrid oligo. **(D)**. DNA loading control (Cp values for amplicons at *ARG4*). Columns: (1) 3 hours with hybrid oligo; (2) 3 hours without hybrid oligo; (3) 4 hours with hybrid oligo; (4) 4 hours without hybrid oligo. **(E).** Intramolecular ligation efficiency calculated as ([intramolecular ligation amplification efficiency]^[-Cp$_{\text{(ligation)}}$]/[*ARG4* amplification efficiency]^[-Cp$_{(ARG4)}$]). Columns as in (D). **(F).** Crosslinking efficiency calculated as ([ssDNA amplification efficiency]^[-Cp$_{\text{(ssDNA)}}$]/[*ARG4* amplification efficiency]^[-Cp$_{(ARG4)}$]). Columns as in (D). **(G).** DNA content at HO cut sites, calculated as ([HO cut site ligation amplification efficiency]^[-Cp$_{(HOcs)}$]/[*ARG4* amplification efficiency]^[-Cp$_{(ARG4)}$]). The "0h" value represents the mean across all available 0-hour time point groups from different genome types with a hybrid oligo added.

### Extended View Figure 8

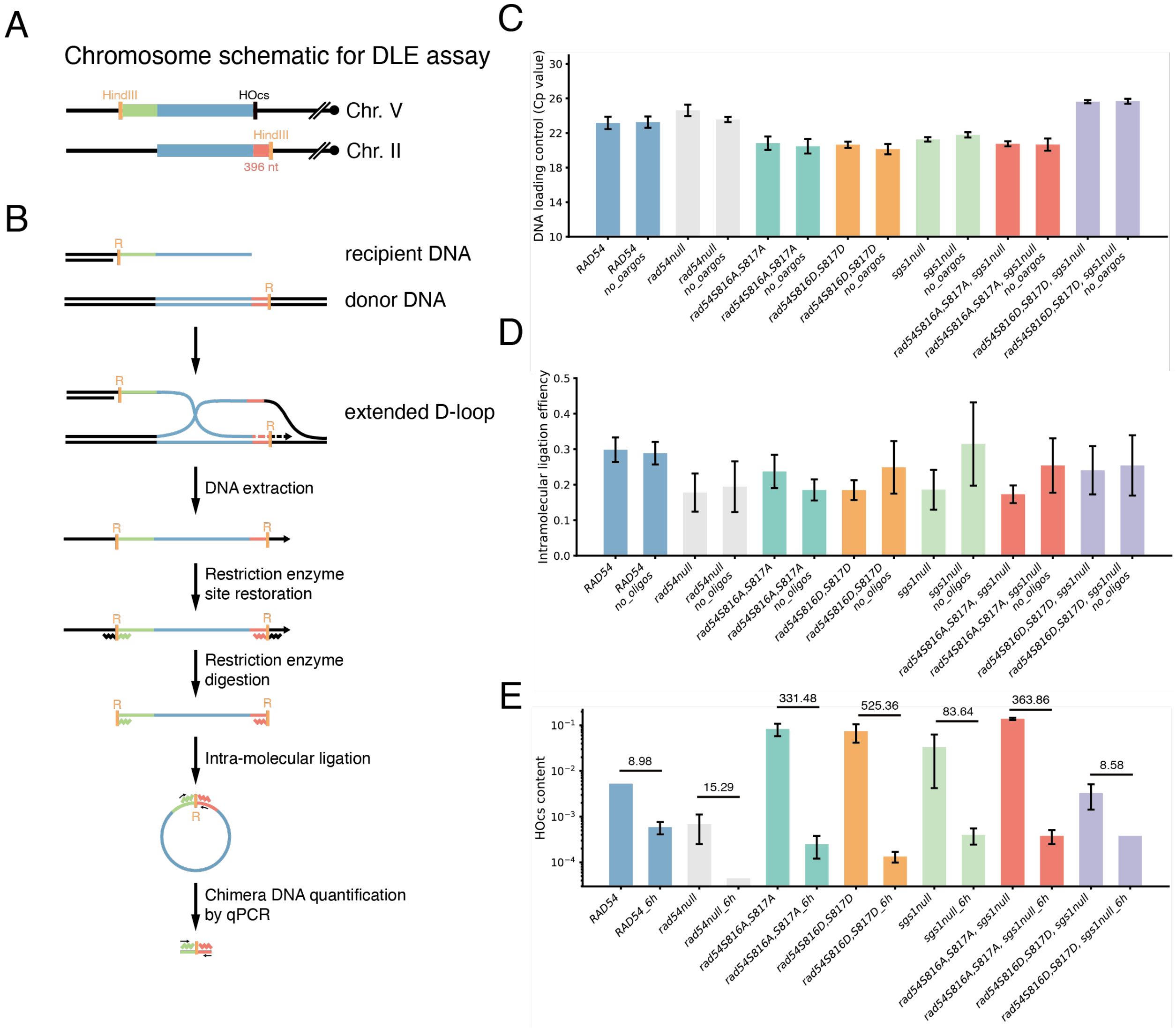


**Extended View Figure 8: Supplementary information for the DLE assay**

**(A).** Chromatin schematic for the DLE assay. **(B).** Rationale for the DLE assay. **(C).** DNA loading control (Cp values for amplicons at *ARG4*). **(D).** Intramolecular ligation efficiency calculated as ([intramolecular ligation amplification efficiency]^[-Cp$_{(ligation)}$]/[*ARG4* amplification efficiency]^[-Cp$_{(ARG4)}$]). **(E).** DNA content at HO cut sites ([HO cut site ligation amplification efficiency]^[-Cp$_{(HOcs)}$]/[*ARG4* amplification efficiency]^[-Cp$_{(ARG4)}$]) at 0-hour and 6-hour time points with hybrid oligos added. Numbers above the bars indicate the reduction of DNA content at the HO cut site after 6 hours.

### Extended View Figure 9

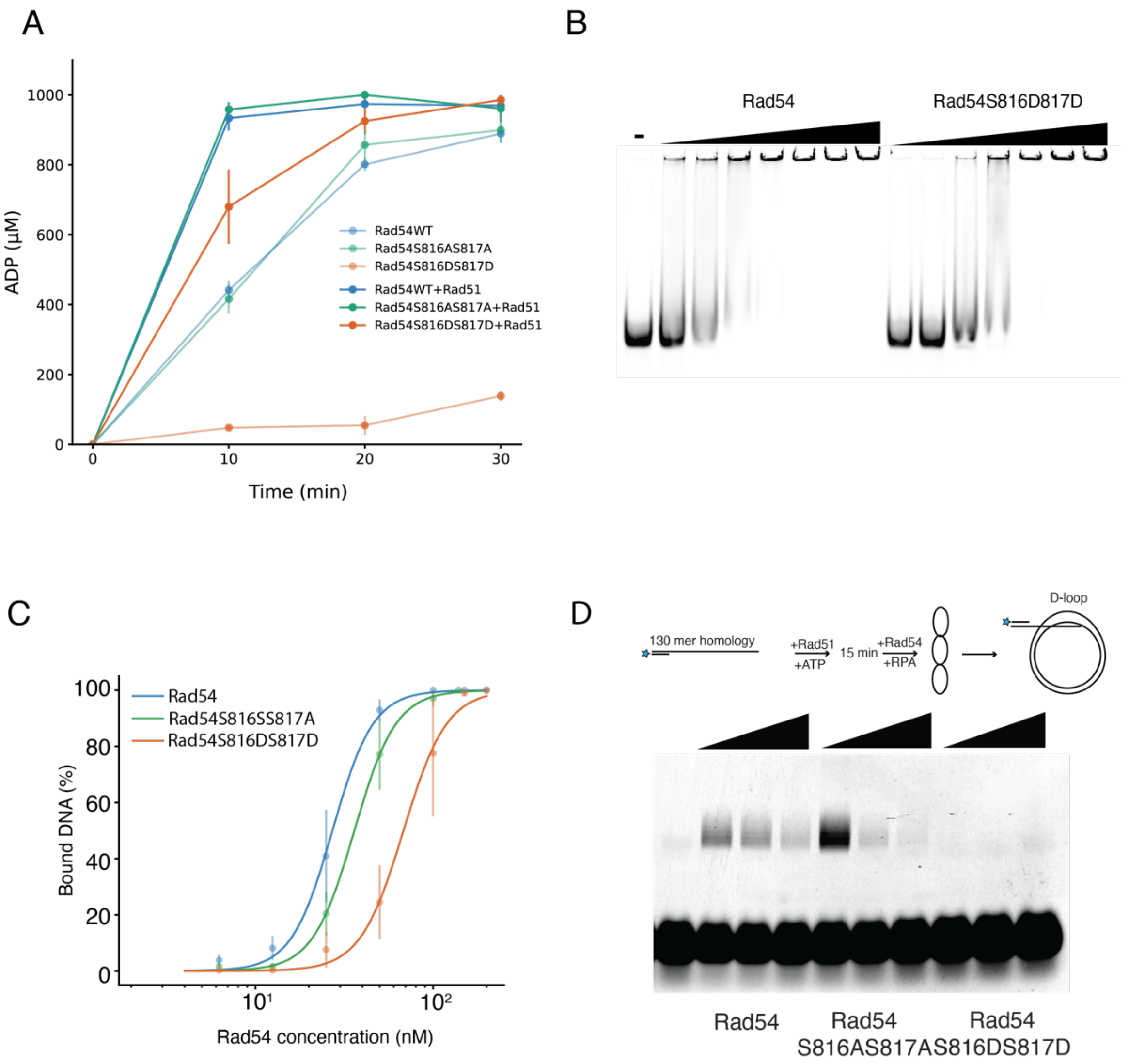


**Extended View Figure 9: Rad54 S816DS817D has lower ATP hydrolysis activity, lower dsDNA binding affinity and lower *in vitro* D-loop formation efficiency.**

**(A).** Line graph representing ATP hydrolysis experiments for Rad54, Rad54 S816A/S817A, and Rad54 S816D/S817D with and without Rad51. The error bars represent the standard error measurement of three independent experiments. **(B).** Representative electromobility shift assay (EMSA) for Rad54 and Rad54 S816DS817D. **(C).** Quantification of EMSA for Rad54, Rad54 S816A/S817A, and Rad54 S816D/S817D is performed using a Hill equation. Fitting is done by neutcurve. The dots represent the mean, and the error bars represent the standard error measurement of the data. **(D).** Representative in vitro D-loop formation experiment for Rad54, Rad54S816A/S817A, and Rad54S816D/S817D.
